# Spatiotemporal WNT and BMP gradients orchestrate regional enteroendocrine cell diversity along the *Drosophila* midgut

**DOI:** 10.1101/2025.08.27.672561

**Authors:** Jiaying Lv, Xingting Guo, Rongwen Xi

## Abstract

Enteroendocrine cells (EEs) in metazoans compromise multiple subtypes that are distributed in specific regions along the digestive tract to perform distinct functions. However, how this regional organization of EE subtypes is established remains poorly understood. Here, we investigated the distribution of three major EE subtypes along the length of the pupal and adult *Drosophila* midgut and found that regional EE pattern is regulated by WNT and BMP signaling in a spatiotemporally dependent manner. Furthermore, both Notch-dependent and -independent modes of cell division by EE progenitors contribute to the generation of regional EE diversity. Our findings suggest that the intercalated WNT and BMP morphogen gradients emanating from compartment boundaries play critical roles not only in establishing regional ISC identity and the resulting EE diversity during development but also in maintaining regional EE diversity in adulthood. This mechanism may represent a conserved paradigm in mammals.

## Introduction

As an important organ for food digestion and nutrient absorption, the intestine is compartmentalized to assign labor along the length of digestive tract for efficient digestion, absorption and metabolic regulation. The spatiotemporal regulation of intrinsic transcription factors (TFs) and extrinsic signaling pathways is involved in the development and specification of intestinal epithelial cells to confer their regional specific identity and function, and abnormal regulation of these factors and pathways may result in intestinal dysfunction and metaplasia ^1–5^.

Enteroendocrine cells (EEs), which originate from local intestinal stem cells (ISCs), exhibit multiple subtypes that are typically located in specific regions of the digestive tract, where they are responsible for sensing stimuli from the gut lumen and secreting peptide hormones and neurotransmitters, facilitating communication with local cells as well as distant organs to regulate various physiological processes ^6,7^. Recent single-cell RNA sequencing (ScRNA-seq) studies have provided a high-resolution understanding of EE heterogeneity in both mammals and *Drosophila* ^8–12^. Additionally, the spatial location of each EE subtype often reflects their regional adaptability to specific biological functions ^6^. For instance, L cells, mainly distributed in the lower small intestine and large intestine, exert negative feedback regulation by secreting glucagon-like peptide-1 (GLP-1), which suppresses appetite and promotes insulin secretion, thereby maintaining plasma glucose homeostasis ^13,14^. GLP-1 agonists have been developed for clinical use to treat diabetes and obesity ^15,16^. Hence, understanding regional EE diversity will not only enhance our knowledge of gut compartmentation and function, but may also provide insight into metabolic diseases and other EE-dysfunction related diseases.

Due to the conservation of gut epithelial lineages and regulatory signaling pathways ^1,6,17,18^, combined with its small size and powerful genetic tools, *Drosophila* offers a compelling genetic model for investigating regional regulation in the digestive tract. The *Drosophila* intestine is divided into the foregut, midgut and hindgut based on its developmental origin. The endodermal-derived midgut encompasses five major anatomical regions (R1-R5) (**Figure 1—figure supplement 1A**) and can be further subdivided into 13 subregions with different metabolic and digestive functions ^19,20^. Similar to mammals, *Drosophila* EEs are also derived from ISCs ^21,22^. A transient activation of the transcription factor Scute induces ISC to generate a transient EE progenitor (EEP), which undergoes one round of asymmetric cell division before terminal differentiation, yielding a pair of EEs ^23,24^. Recent single-cell analyses of EEs in adult *Drosophila* midgut have led to the identification of totally 10 EE subtypes, which can be generally categorized into three major classes ^12,25^. These EE subtypes are dispersed throughout the adult *Drosophila* midgut and also exhibit specific regional distribution and functional adaptation ^26–30^. For instances, Neuropeptide F (NPF)-secreting EEs residing in the anterior and middle midgut can sense sugar intake to regulate glucose and lipid metabolism ^31^; Allatostatin C (AstC)-expressing EEs enriched at the posterior midgut can be activated by starvation to promote food intake and energy mobilization ^32^; and Diuretic hormone 31 (DH31)-expressing EEs are located nearby Malpighian tubule and have been implicated in regulating water flux homeostasis ^18^. The mechanisms driving these distribution patterns remain unclear.

One potential mechanism underlying regional EE diversity is the regional identity of ISCs. ISCs located in specific regions of the gut possess distinct regional identities and therefore give rise to region-specific progeny. A well-studied example is the development of the copper cell region (CCR) in the middle midgut, where BMP signaling is activated to induce *lab* expression. Consequently, it promotes the adoption of a gastric stem cell identity, enabling these ISCs to generate acid-secreting copper cells rather than enterocytes ^33,34^. However, ISCs in the anterior and posterior midgut exhibit largely similar transcriptomic profiles and cluster together in single-cell principal component analysis (PCA) ^25^. Furthermore, no specific gene markers have yet been identified that distinguish ISCs in the anterior midgut from those in the posterior region. These findings suggest that additional mechanisms may contribute to the regulation of regional EE diversity.

In this work, we characterized the regional distribution of EE subtypes (at the class level) and asymmetric cell division patterns of EEPs along the length of pupal and adult *Drosophila* midgut, and investigated the function of the extrinsic niche signals, including Notch, WNT and BMP pathways, in the regulation of regional EE diversity. Our results suggest the participation of both intrinsic (regional ISC identity) and extrinsic (morphogen gradient) mechanisms in shaping regional EE diversity.

## Results

### The distribution patterns of class I, II and III EEs along the length of the *Drosophila* midgut

In addition to AstC^+^ class I and Tachykinin^+^ (Tk) class II EEs, our recent scRNA-seq analysis of the EEs has revealed the existence of another EE subtype named as class III EEs, which express neither AstC nor Tk, but do express peptide hormone CCHamide-2 (CCHa2) ^12^ (**Figure 1—figure supplement 1A**). We confirmed this pattern in the R2 region using a Gal4 knock-in line of CCHa2 (**Figure 1—figure supplement 1B**). Interestingly, CCHa2 was also found to be expressed in a subset of AstC^+^ class I EEs in the R4 region (**Figure 1—figure supplement 1C-E**). Therefore, class III EEs are localized to the R2 region with a specific expression pattern of peptide hormones: CCHa2^+^, AstC^-^, Tk^-^.

Combining the region-specific GAL4 enhancer trap lines ^19^ with the immunostaining of CCHa2 antibodies, we found that CCHa2^+^ class III EEs began to appear in the R1b region and disappeared at the junction between R2 and R3 regions (**Figure 1—figure supplement 1F and H**). CCHa2 expression reappeared in the R4b and R4c regions and disappeared in the R5 region (**Figure 1— figure supplement 1G-H**). Consistent with previous studies ^12^, Tk^+^ class II EEs are distributed along the whole gut except for the R4b region, where EEs are all belong to AstC^+^ class I (**Figure 1—figure supplement 1G-H**).

Based on this information, we next co-stained AstC, Tk and Pros (Prospero, a pan-EE marker) in the midgut of *CCHa2-LexA, LexAOP-GFP* (abbreviated as *CCHa2-LexA>GFP*) flies to determine the distribution pattern of class I, II and III EEs along the length of midgut at subregion resolution (**Figure 1**). The AstC^+^ class I and Tk^+^ class II EEs were found in the R3-R4a and R4c-R5 regions with the ratio of approximately 1:1 (**Figure 1D, 1F and 1G**), a pattern that is consistent with the traditional view that each EEP divides asymmetrically prior to differentiation, yielding one class I EE and one class II EE ^23,35^. The rest of regions or subregions, however, do not exactly follow this pattern.

**Figure 1.**
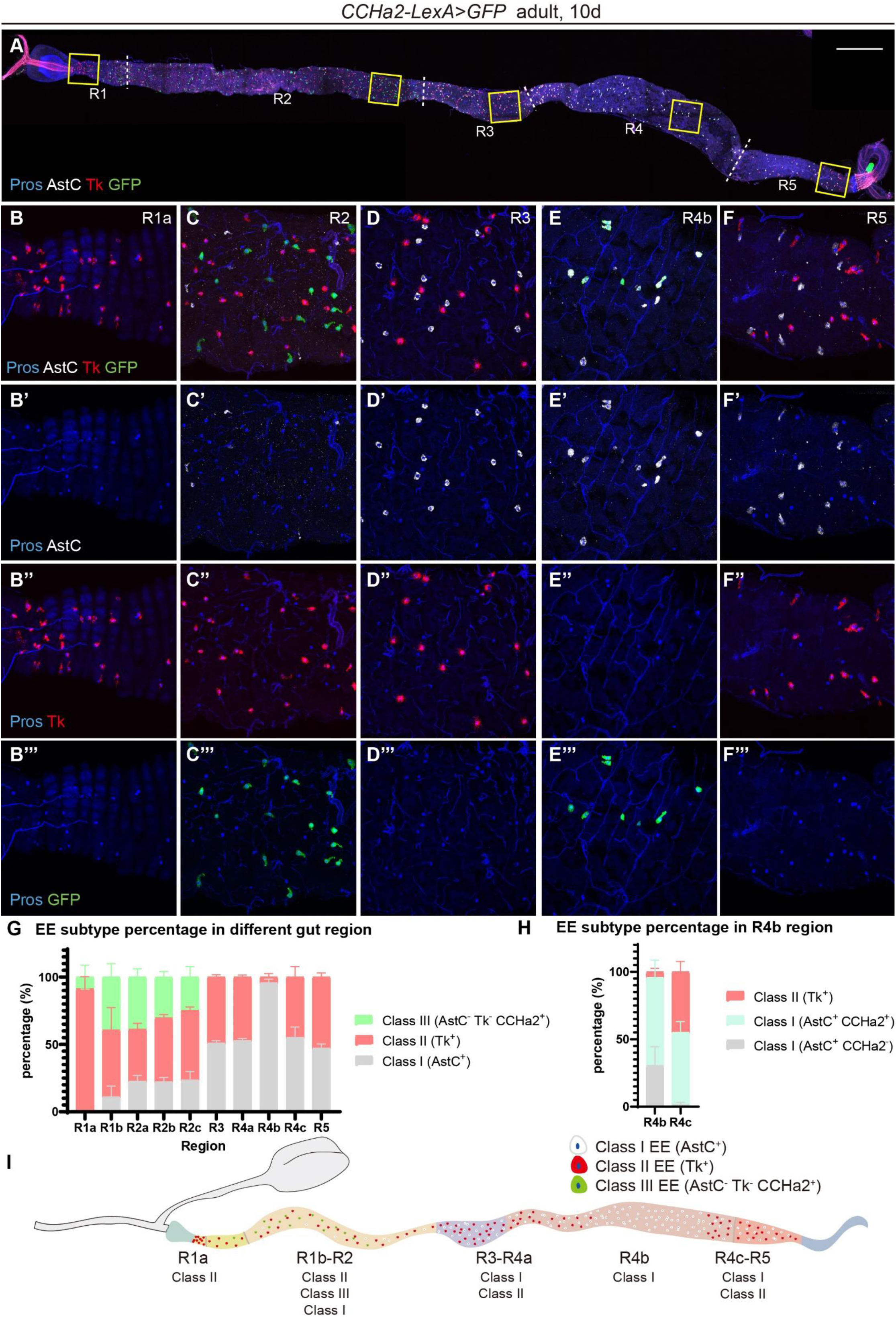
The distribution pattern of EE subtypes along the length of adult Drosophila midgut. (A-F) Midguts of *CCHa2-LexA>GFP* flies stained with anti-Pros (blue), anti-AstC (white) and anti-Tk (red) antibodies 10 days after eclosion. Separated color channels for the insets from different gut regions in (A) were enlarged and displayed in (B-F), showing EE distribution characteristic of different gut regions. Scale bar represents 300 μm. (G) Quantification of the three major EE subtype percentage in different gut regions of flies 10 days after eclosion. Error bars represent Mean ± SEM, n=5. (H) Quantification of EE subtype percentage in the R4bc region of flies 10 days after eclosion. Error bars represent Mean ± SEM, n=5. (I) A schematic diagram showing the distribution patterns of the three EE subtypes along the *Drosophila* midgut.

In the R1a region, the resident EEs are almost all Tk^+^ class II EEs (**Figure 1B and 1G**); The R1b-R2 region is rather complex as all three classes of EE subtypes are found in this area: The class II EEs accounts for about half of the total EE number in this region, followed by class III EEs, accounting for about 30-40%, and the rest are class I EEs (**Figure 1C and 1G**). Moreover, the proportions of three EE subtypes in R1-R2 change in a gradient manner, with more class III EEs towards anterior, and more class I and II EEs towards posterior (**Figure 1G**).

The R4b region is even more surprising, as virtually all EEs in this region are AstC^+^ class I EEs (**Figure 1E and 1G**). In addition, a subset (about three-quarters) of these EEs is CCHa2^+^ (**Figure 1H**). The annotated distribution pattern of class I, II and III along the length of midgut is shown in **Figure 1I**. The distinct regional EE patterns, especially found in R1a, R1b-R2 and R4b provide an opportunity for the genetic dissection of the mechanisms underlying regional EE diversity.

### The EE distribution pattern is established at late pupal stage

During the pupation of *Drosophila*, the midgut epithelium undergoes dramatic changes to produce definitive adult midgut ^36,37^. Specified prior to pupal formation, EEs are lost at early pupal stage, but re-emerge later from pupal ISCs ^36,38^. Therefore, we wondered whether the regional diversity of EE subtypes was established at this stage.

Similar to the adult midgut, we found that the regional boundary markers *GMR46B08-Gal4* and *GMR50A12-Gal4* have already expressed in the midguts at late pupal stage— 96h after pupa formation (APF) (**Figure 1—figure supplement 1I-J**). Although the expression domain was wider than that in adult midguts (**Figure 1—figure supplement 1G-H**), the trends of regional-specific expression had already been determined. Immunostaining of pupal midguts with Tk, AstC and CCHa2 showed that the distribution pattern and proportion of class I, II and III EE subtypes in different regions were virtually the same as those in adults (**Figure 1—figure supplement 2**). We propose that *Drosophila* midgut compartmentation and regional EE subtype diversity were established at the pupal stage and maintained to the adulthood. After pupal formation, the pupal ISCs undergo two rounds of cell divisions to generate EE mother cells or EEPs, which then divide once to yield a pair of EEs ^24,36,38^. Therefore, EEs found in late pupal midgut are all generated by pupal ISCs, implying that regional EE diversity may be determined or contributed by the regional identity of ISCs.

It is worth mentioning that in the R5 region of the late pupal midgut, approximately half of the Pros^+^ EEs were Tk^+^ class II, but the other half were negative for AstC expression, so as for CCHa2 (**Figure 1—figure supplement 2F and H**). Considering the expression pattern in the newly enclosed and adult flies, in which these cells are positive for AstC (**Figure 1F-G and Figure 1—figure supplement 2G**), these R5 EEs at the pupal stage are likely immature EEs, which are expected to reach maturity shortly after eclosion.

### Both symmetric and asymmetric division of EEPs contribute to regional EE diversity

In the adult midgut, EEPs undergo a single asymmetric division to produce a pair of EEs, typically consisting of one class I EE and one class II EE ^23,35^. Given the observations that several gut regions, including R1, R2 and R4b, do not harbor this typical 1:1 EE pattern, we asked how different subtypes of EEs are specified from EEPs. Because ISCs are normally quiescent and EEPs are rarely found in the intestinal epithelium, we induced intestinal damage by feeding flies with 5% Dextran Sulfate Sodium (DSS) solution ^39^, which can rapidly induce ISC proliferation and differentiation to replenish damaged enterocytes and EEs. Then the newly generated EE pairs can be distinguished by two EEs that are in close proximity with each other ^23,35^ (**Figure 2A-E**). As expected, the majority of the EE pairs found in the R3-R4a and R4c-R5 regions consisted of one class I and one class II EE (**Figure 2C and 2E-F**), which fits into the canonical view that each EEP divides asymmetrically to yield class II/I EE pair.

**Figure 2.**
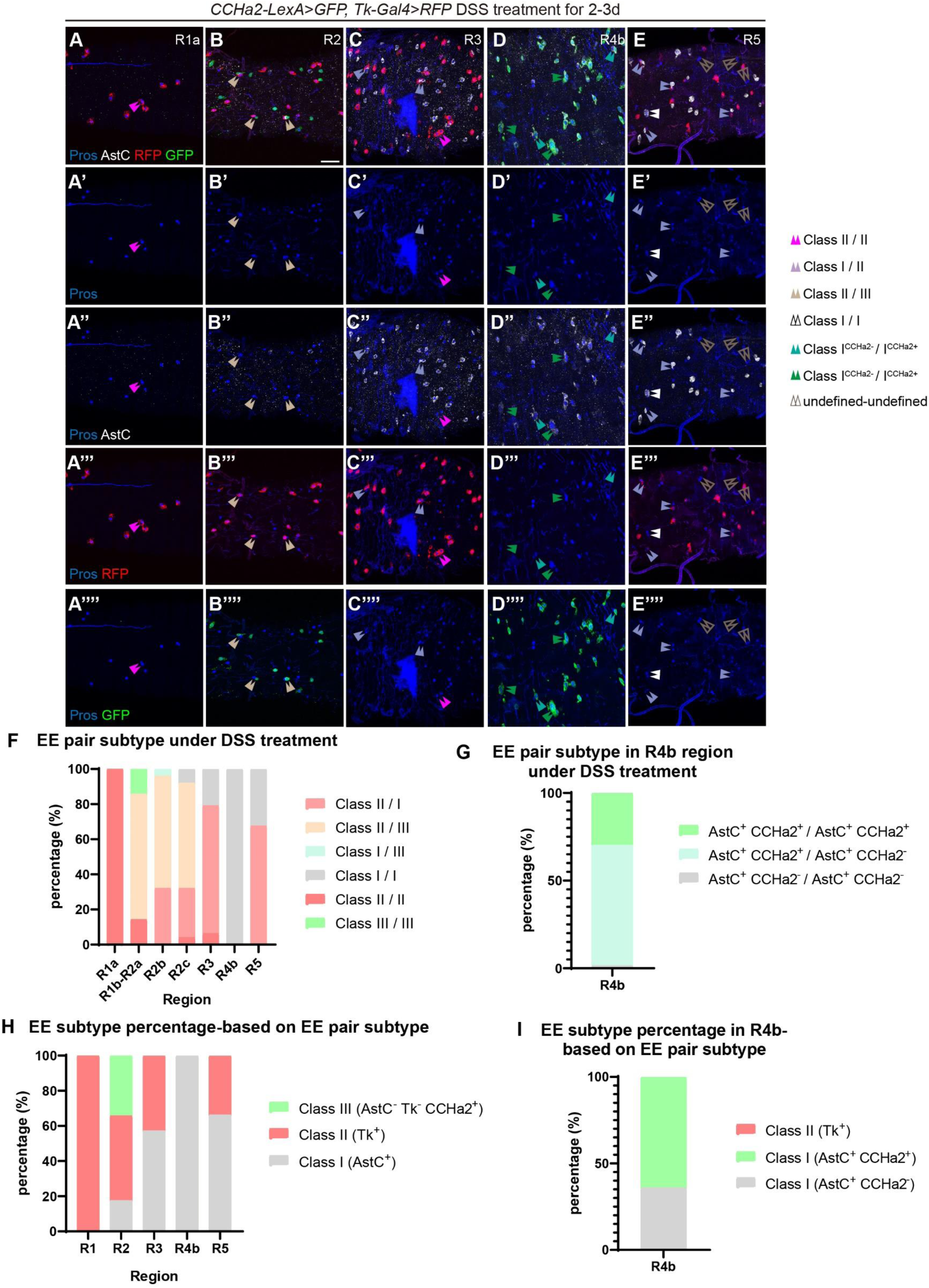
Choices of symmetric or asymmetric division of EEPs contribute to regional EE diversity. (A-E) Midguts stained with anti-Pros (blue) and anti-AstC (white) antibodies in *CCHa2-LexA>GFP, Tk-Gal4>RFP* flies fed with 5% DSS for 2-3 days. EE pairs were pointed by arrow head pairs. Scale bar represents 25 μm. (F) Quantification of different EE subtype pair percentage in different gut regions. (R1a: n=14; R1b-R2a: n=7; R2b: n=25; R2c: n=25; R3-R4c: n=48; R4b: n=64; R4c-R5: n=71) (G) Quantification of different EE subtype pair percentage in the R4b region. (n=64) (H) Speculation of EE subtype percentage in different gut regions by taking the percentage of EE subtype pairs from (F) to generate EEs. (I) Speculation of EE subtype percentage in the R4b region by taking the percentage of EE subtype pairs from (G) to generate EEs.

However, the two EEs in the EE pairs found in the R1a region were all Tk^+^ (**Figure 2A and 2F**), and in the R4b region were all AstC^+^ (**Figure 2D and 2F**), suggesting that EEPs in these two regions undergo symmetric cell division to yield identical or “symmetric” EE pairs.

We next examined the EE pairs in R1b-R2 regions, where class I, II and III EE subtypes co-reside. We found that the majority of EE pairs were asymmetric pairs, as about 60% EE pairs consist of Tk^+^ class II + CCHa2^+^ class III EE, and 35% consist of Tk^+^ class II + AstC^+^ class I EE (**Figure 2B and 2F**). The remaining symmetric pairs or asymmetric class I/III pairs were rarely found. The percentage of these asymmetric pairs were largely consistent with the percentage of class I, II and III EE subtypes found in these regions (**Figure 2H and 1G**), suggesting that EEs in R1b-R2 regions are generated through these two modes of EEP asymmetric cell divisions, one that generates class II/III EE pair, and the other that generates class II/I EE pair.

It worth mentioning that, although the EE pairs found in R4b region were considered symmetric AstC^+^/AstC^+^ class I EE pair (**Figure 2D and 2F**), we found that about 70% of these EE pairs were not strictly identical, with one expressing CCHa2 while the other not (**Figure 2D and 2G**), and the consequence of these cell division modes were largely consistent with the percentage of EE subtypes found in this region (**Figure 2I and 1H**). How this heterogeneity occurs is unclear but as described shortly, the loss of Notch does not affect this CCHa2 pattern in the AstC^+^ EE pairs, suggesting this heterogeneity is not generated by Notch-mediated lateral inhibition.

It is known that tissue damage caused by DSS treatment can induce the activation of multiple signaling pathways from the ISC environment, which could potentially affect the regular generation of regional EE diversity. We therefore also used a different approach to induce EE pair formation. Overexpression of *scute* in ISCs could induce rapid ISC proliferation and differentiation toward EE lineage and form EE clusters ^23,40^. We therefore conditionally induced *scute* overexpression in ISCs for 12-48 hours, which led to a large number of EE pairs (**Figure 2—figure supplement 1A-E**). We then performed a similar analysis and quantified the proportion of symmetric or asymmetric EE pairs in each gut regions, the results were largely consistent with that obtained under DSS treatment conditions (**Figure 2—figure supplement 1F-I**). Therefore, symmetric and asymmetric division of EEPs varies across different regions of the gut, thereby contributing to regional EE subtype diversity.

### Notch is required for the specification of class II but not for class I and III EEs

It is known that Notch is required for the asymmetric division of EEPs to generate asymmetric class I/II EE pairs, as the loss of Notch signaling leads to the loss of class II EE lineage ^35,41^. But it is unclear if Notch is required for the specification of class III EEs, and for the generation of II/III and class I/III asymmetric EE pairs found in R2 region.

We used MARCM system ^42^ to generate Notch knockdown or Dl mutant clones. Compared to WT clones (**Figure 3—figure supplement 1A and C**), all Notch RNAi or *Delta^Revf^*^10^ (abbreviated as *Dl^Revf^*^10^) homozygous clones in the R2 region consist of clustered small diploid cells that include CCHa2^+^ class III and AstC^+^ class I EEs, while class II EEs were only observed outside of the mutant clones (**Figure 3—figure supplement 1B and D**). These results suggest that local Delta-Notch signaling is essential for the establishment of class II but not class I or III EEs.

To further validate the role of Delta-Notch signaling in EE subtype differentiation along the whole midgut, we introduced Notch RNAi in ISCs for 5-10 days in adult midguts, which led to the generation of many EE clusters (or EE tumors derived from Notch-depleted ISCs) along the length of the midgut (**Figure 3**). We then examined the EE subtype identity in these EE clusters in R1-R5 regions respectively. Interestingly, no EE clusters were found in R1a (**Figure 3A**), indicating that ISCs in this region are relatively quiescent. As expected, in R3-R4a and R4c-R5, where normally only class I/II EEs reside, all EEs in the EE clusters became AstC^+^ class I EEs (**Figure 3C and 3E**).

**Figure 3.**
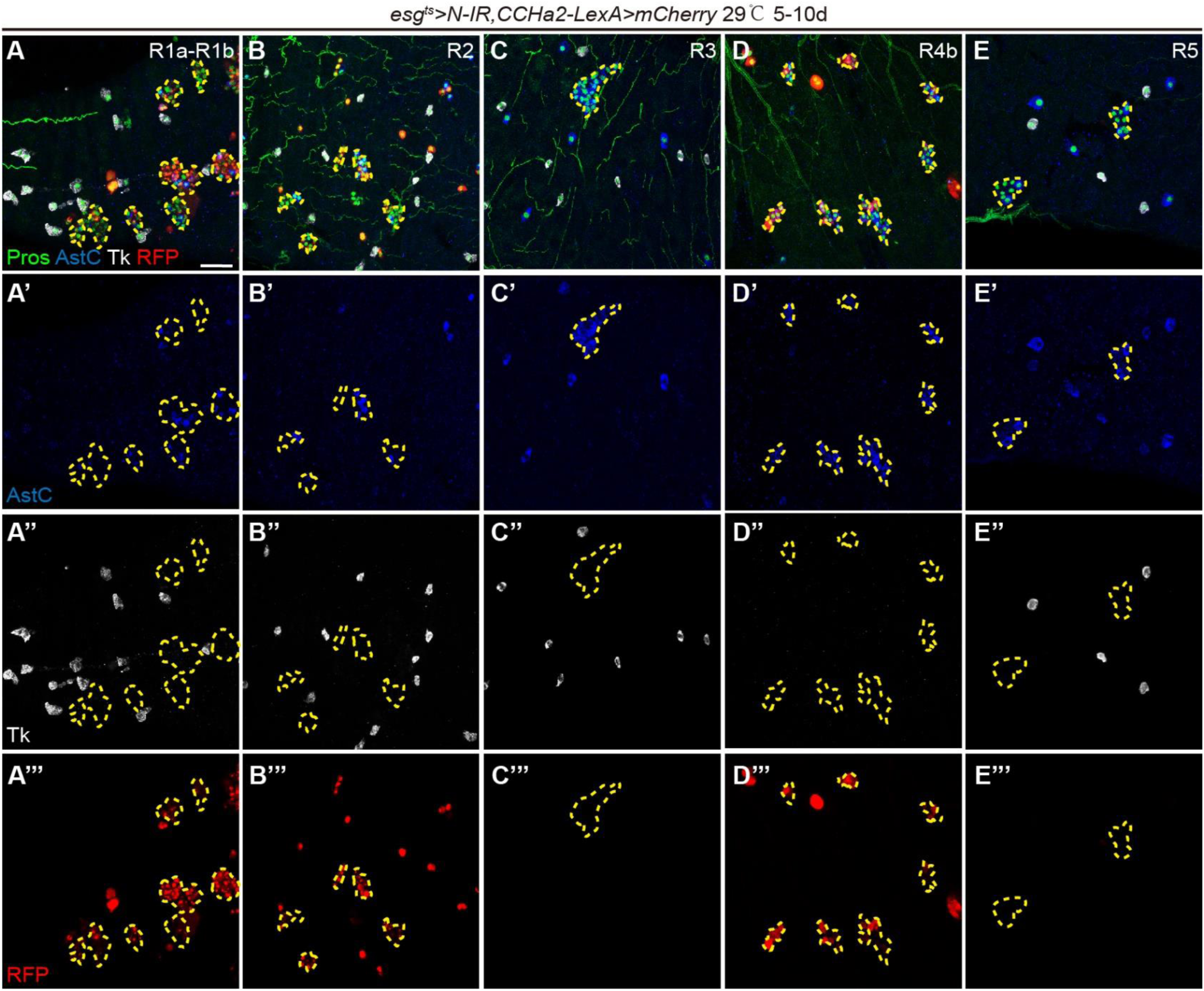
Notch is dispensable for the specification of both class I and III EE subtypes. (A-E) Midguts stained with anti-Pros (green), anti-AstC (blue) and anti-Tk (white) antibodies in *esg-Gal4^ts^>Notch RNAi, CCHa2-LexA>mCherry* flies shift to 29 ℃ for 5-10 days after eclosion. Scale bar represents 25 μm.

In R1b-R2, only AstC^+^ and CCHa2^+^ EEs, but not Tk^+^ EEs were found in the EE clusters (**Figure 3A-B**), showing that the specification of class II EE is failed in these regions. For most EE clusters, we found that AstC^+^ EEs and CCHa2^+^ EEs were equally presented and mixed together in a salt and pepper fashion. Clusters containing more class III EEs and rare class I EEs were observed occasionally, but clusters containing only class I EEs were never observed (**Figure 3A-B**). To further assess whether class I and class III EE specification occur independently of Notch activity, we quantified their relative proportions within EE clusters. On average, class I and class III EEs accounted for approximately 30% and 70%, respectively (**Figure 3—figure supplement 2A**), a distribution consistent with that observed in wild-type anterior midguts (**Figure 1G**). Based on these observations, we hypothesize that loss of Notch does not abolish asymmetric division of EEPs in the R1b-R2 regions but instead alters the resulting cell fate outcomes. Specifically, it may shift EE pair composition from class II/I or II/III toward class III/I or III/III, without preferentially favoring either class III or class I EEs.

In R4b, the EE clusters generated by Notch-depleted ISCs remained as AstC^+^ EE clusters, and approximately 65% of these EEs were also positive for CCHa2 expression, a proportion consistent with that observed in wild-type midguts (**Figure 1H, 3D and Figure 3—figure supplement 2B**). Thus, the asymmetric pattern of CCHa2 expression appears to be remained despite the loss of Notch activity. Nevertheless, given that class III EE production remains spatially restricted to the R1b–R2 regions, whereas CCHa2 continues to be co-expressed with AstC in the R4 region, additional regulatory factors likely act in concert with Notch to establish regional identity by specifying class III EE fate and regulating asymmetric cell division.

Taken together, these observations suggest that Notch is required for class II EE specification but not for class I nor III EEs, and it appears that Notch is not involved in the asymmetric cell division of EEPs to yield class I/III EE pairs in R2, nor in the asymmetric expression of CCHa2 within symmetric class I/I EE pairs in R4b.

### Wnt morphogen gradient orchestrates regional patterning of class II EEs

We next investigated how the specific regional patterns of EE subtypes are established from pupal ISCs, especially in R1a, R1b-R2 and R4b regions. Given the gradual changes in the proportion of the three EE subtypes found in R1-R2 region (**Figure 1G**), we speculated that signaling gradients, such as WNT gradient, emanated from gut compartment boundaries, might be involved. It has been shown that the WNT ligand Wingless (Wg), which is expressed in the visceral muscle and the epithelial cells at several junctional regions, including the proventriculus-R1, R2-R3, and R5-hindgut boundaries, generates WNT signaling gradient and is important for the development of normal gut morphology and regulation of ISC proliferation ^43–45^. Consistent with the WNT gradient formation at the junctional regions, we found that WNT4 showed a similar expression pattern to Wg reflected by a GAL4 enhancer trap line (**Figure 4—figure supplement 1A**), and it is also similar for Wnt6 (data not shown). Moreover, we also found that in both pupal and adult midgut, *Fz3-RFP,* a WNT activation reporter, showed a gradient pattern in the epithelium emanated from the junctions (**Figure 4A and Figure 4—figure supplement 1B-F**). Its expression in adult ISCs is observed throughout the entire midgut and also displays a gradient that varies across different gut regions (**Figure 4—figure supplement 1G**).

**Figure 4.**
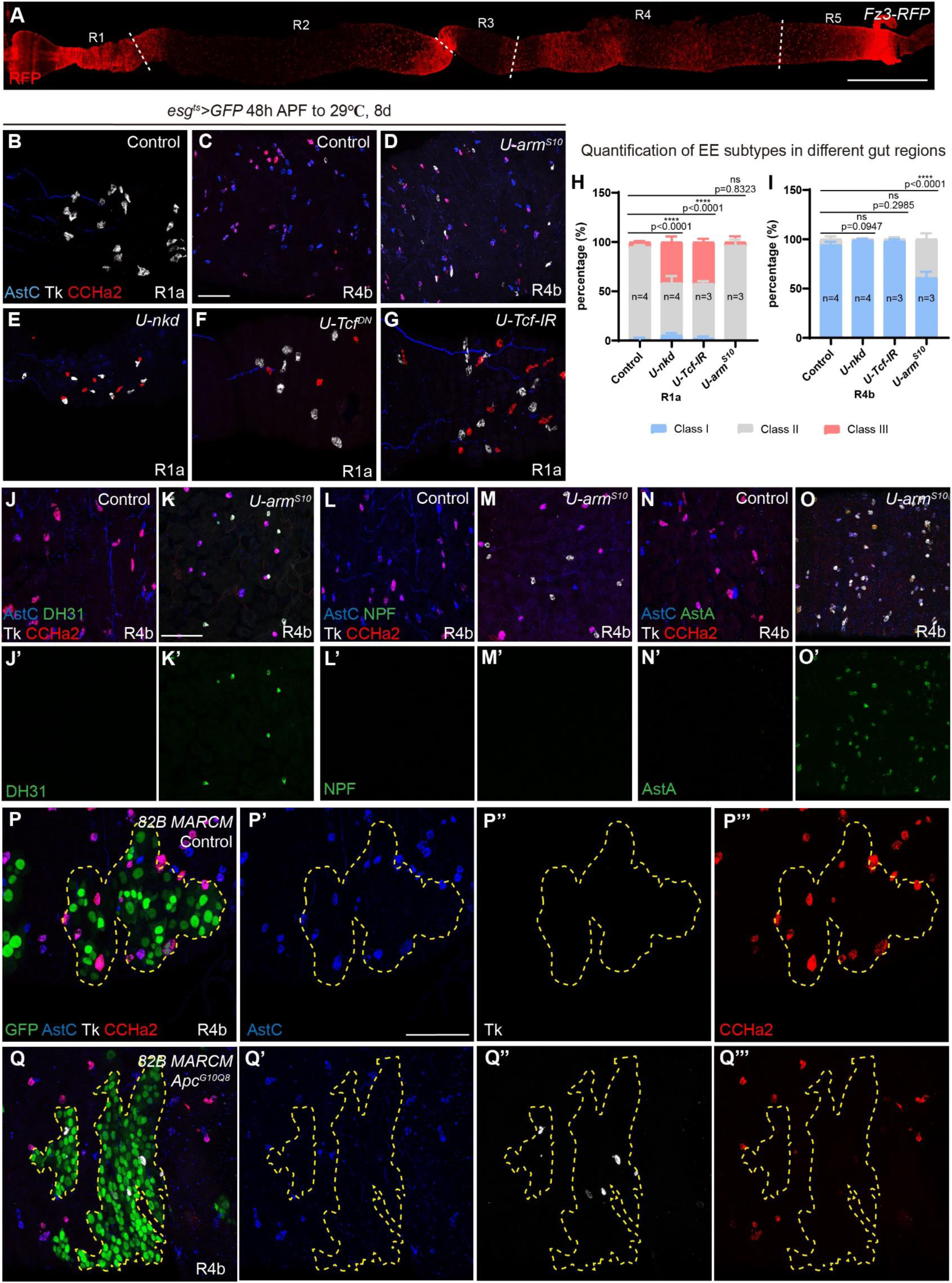
WNT signaling gradient orchestrates regional class II EE specification. (A) Midguts of *Fz3-RFP* flies stained with anti-RFP (red) antibodies 5 days after eclosion. (B-G) Midguts stained with anti-AstC (blue), anti-Tk (white) and anti-CCHa2 (red) in the following genotypes of flies driven by *esg-Gal4^ts^* for 8 days since 48h APF: Control (B-C), *UAS-arm^S^*^10^ (D), *UAS-nkd* (E), *UAS-Tcf^DN^* (F) and *UAS-Tcf RNAi* (G). (H-I) Quantification of the three major EE subtype percentage in R1a (H) and R4b (I) of Control, *UAS-nkd*, *UAS-Tcf RNAi* and *UAS-arm^S^*^10^ flies. Error bars represent Mean ± SEM, ****p <0.0001 (two-way ANOVA). (J-K) Midguts stained with anti-AstC (blue), anti-DH31 (green), anti-CCHa2 (red) and anti-Tk (white) in the following genotypes of flies driven by *esg-Gal4^ts^* for 10 days since 48h APF: Control (J-J’) and *UAS-arm^S^*^10^ (K-K’). (L-M) Midguts stained with anti-AstC (blue), anti-NPF (green), anti-CCHa2 (red) and anti-Tk (white) in the following genotypes of flies driven by *esg-Gal4^ts^* for 10 days since 48h APF: Control (L-L’) and *UAS-arm^S^*^10^ (M-M’). (N-O) Midguts stained with anti-AstC (blue), anti-AstA (green), anti-CCHa2 (red) and anti-Tk (white) in the following genotypes of flies driven by *esg-Gal4^ts^* for 10 days since 48h APF: Control (N-N’) and *UAS-arm^S^*^10^ (O-O’). (P-Q) MARCM clones co-stained with anti-AstC (blue), anti-Tk (white) and anti-CCHa2 (red). (P-P’’’) WT *FRT82B* clone, (Q-Q’’’) *Apc^G10Q^*^8^ *FRT82B* MARCM. Scale bar in (A): 300 μm; others: 50 μm.

To test if the WNT gradient is important for shaping regional pupal ISC identity and further influence the generation of the observed regional EE subtype patterns, we overexpressed a WNT pathway negative regulator *nkd*, or a dominant negative form of a WNT pathway effector Tcf (*Tcf^DN^*), or RNAi of Tcf (*Tcf-IR*) in ISCs, started since early pupae stage (48h APF), and then examined the distribution pattern of class I, II and III EEs in the midgut 5 days after eclosion. We observed a consistent, modest reduction in *Fz3-RFP* expression in the R1 region across all experimental conditions (**Figure 4—figure supplement 2F-G**). This reduction was associated with the ectopic appearance of class III EEs in the R1a region (**Figure 4B, 4E-G and Figure 4—figure supplement 2A-D**) where they accounted for approximately 45% of the total EE population, a proportion comparable to that normally observed in the R1b and R2 regions (**Figure 4H**). It suggests that a precise high level of WNT signaling activation may characterize the pupal ISC identity specific to the R1a region and facilitate the symmetric division of EEPs, thereby contributing to the formation of class II EE pairs in this region. Conversely, inhibition of WNT signaling may reduce its activity to a level comparable to that observed in the R1b-R2 regions, thereby facilitating asymmetric division of EEPs and resulting in the formation of class II/III EE pairs. Although *Fz3-RFP* expression was also perturbed in R2c and R4a (**Figure 4—figure supplement 2F-G**), regional EE patterning remained largely intact in these regions (**Figure 4—figure supplement 2I-N**), implying that, additional, region-specific factors contribute to the establishment and maintenance of regional ISC identity.

We next examined the effects of WNT signaling activation on class II EE fate specification and regional EE subtype patterns by overexpressed an active form of WNT signaling effector arm (*arm^S^*^10^) (**Figure 4—figure supplement 2E**). The expression of *Fz3-RFP* reporter was upregulated in the middle and posterior midgut (**Figure 4—figure supplement 2H**). We found that although the pattern remained largely normal in anterior and middle midgut (**Figure 4—figure supplement 2I-L**), the proportion of Tk^+^ class II EEs in posterior midgut were increased (**Figure 4—figure supplement 2M-N**), and ectopically appeared in the R4b region (**Figure 4D and 4I**), where they should be absent (**Figure 4C**). Normally class II EEs in R4-R5 regions can typically be distinguished from these in R1-R2 regions by their co-expression of the peptide hormone DH31, and class II EEs in R2-R3 region can typically be distinguished from those in other regions by their co-expression of the peptide hormone NPF ^12^. Virtually all of these ectopically appeared class II EEs in R4b co-expressed peptide hormone DH31 (**Figure 4J-K**) but not NPF (**Figure 4L-M**), suggesting that these EEs are not as a result of class II EE expansion from anterior and middle regions, but represent ectopic formation of class II EEs that similar to those in posterior regions. In addition, we found that the class I EEs in this region ectopically expressed peptide hormone Allatostatin A (AstA) (**Figure 4N-O**), which are normally expressed specifically in the R4c-R5 but not in R4b or other region or subregions ^12^. Taken together, these observations suggest that overactivation of WNT signaling causes the regional EE pattern of R4b region to resemble that of R4c-R5 region.

To further confirm the role of the WNT signaling pathway in the regulation of EE subtype fate, we used MARCM system to generate *Apc^Q^*^8^ and *Apc2^g^*^10^ double mutant clones (abbreviated as *Apc^g10Q^*^8^) in which WNT signaling is hyperactivated ^46^. We found that Tk^+^ class II EEs specifically appeared in *Apc^g10Q^*^8^ homozygous clones, but not the wild type control clones in the R4b region (**Figure 4P-Q**), which further supports the notion that hyperactivation of WNT signaling activity is sufficient to promote the generation of class II EEs in R4b region, where these EEs are normally absent.

Therefore, it appears that a high level of WNT signaling favors the specification of Tk^+^ class II EE identity from pupal ISCs. Inhibition of WNT signaling results in the regional EE patterns of R1a region resembling those of R1b-R2, whereas overactivation of WNT signaling leads to the EE patterns of R4b region adopting characteristics typical of R4c-R5. However, suppression of WNT signaling does not produce fully reciprocal phenotypes in R4c-R5 region. One possible explanation is that additional regulatory mechanisms may cooperate with WNT signaling to modulate regional ISC identity and cell division mode.

### BMP signaling negatively regulates class III EE specification and promotes class I/II EE pair mode

In addition to WNT, BMP is also recognized as an important morphogenetic factor. BMP-like Dpp is highly enriched at the boundary between posterior midgut and middle midgut in *Drosophila* and plays an essential role in the determination of R3 or copper cell region ^33,34,47,48^. Based on the expression of *Dad-LacZ*, a BMP signaling activity reporter, the highest activity was found in the nuclei of muscle cells and epithelial cells within the R3 region at pupal stage (**Figure 5—figure supplement 1A-C**). This activity was also seen both in R3 and R5 at the adult stage (**Figure 5A and Figure 5—figure supplement 1D**), displaying a gradient pattern in the epithelium that emanates from these regions (**Figure 5—figure supplement 1E**). We therefore examined whether BMP signaling has a role in regulating regional EE patterns.

**Figure 5.**
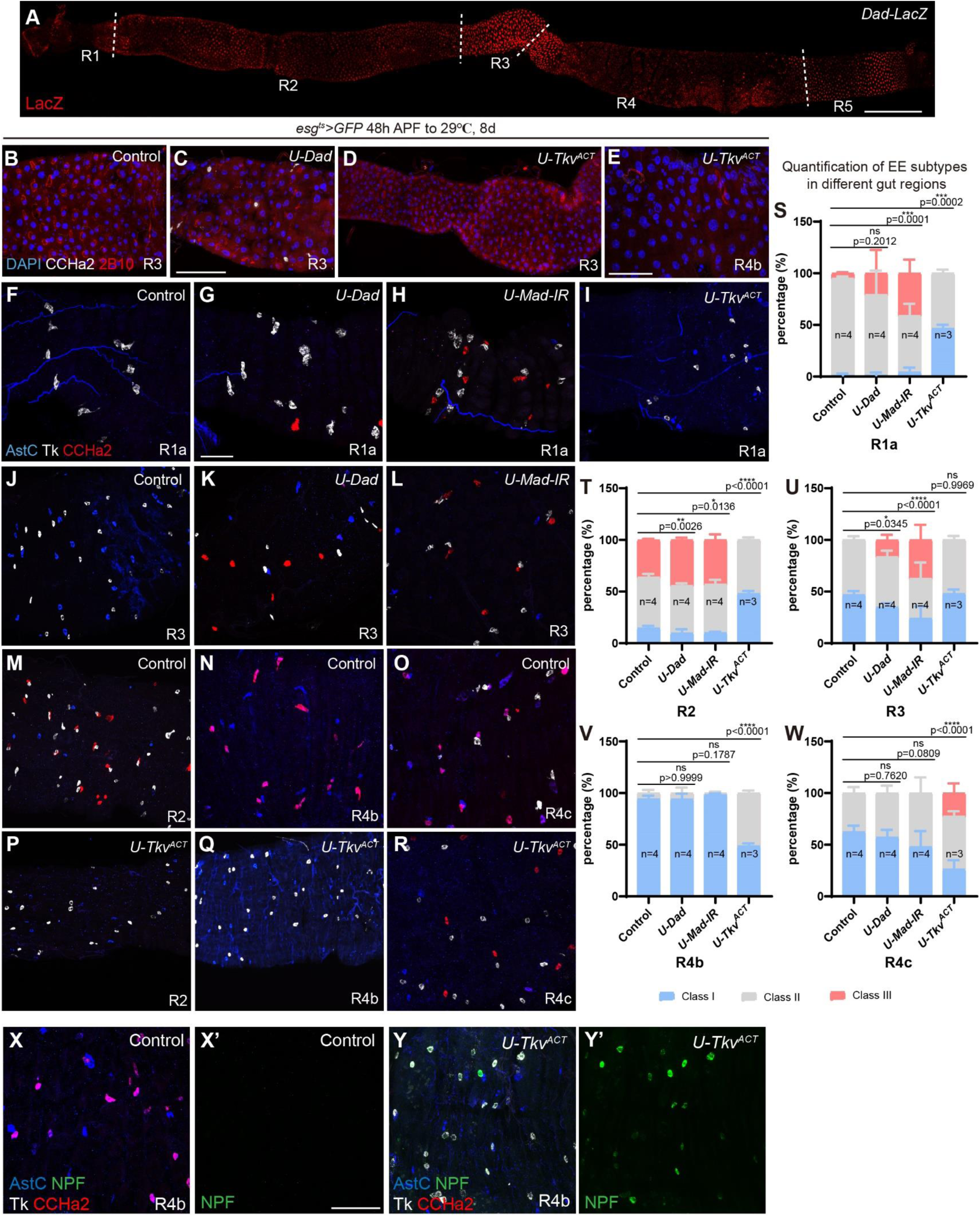
BMP signaling negatively regulate class III EE differentiation. (A) Midguts of *Dad-LacZ* flies stained with anti-LacZ (red) antibodies before eclosion. (B-E) Midguts stained with DAPI (blue), anti-2B10 (red) and anti-CCHa2 (white) in the following genotypes of flies driven by *esg-Gal4^ts^* for 8 days since 48h APF: Control (B), *U-Dad* (C), *U-Tkv^ACT^* (D-E). (F-R) Midguts stained with anti-AstC (blue), anti-Tk (white) and anti-CCHa2 (red) in the following genotypes of flies driven by *esg-Gal4^ts^* for 8 days since 48h APF: Control (F, J and M-O), *UAS-Dad* (G and K), *UAS-Mad RNAi* (H and L) and *UAS-Tkv^ACT^* (I and P-R). (S-W) Quantification of the three major EE subtype percentage in R1a (S), R2 (T), R3 (U), R4b (V) and R4c (W) of *UAS-Dad*, *UAS-Mad RNAi* and *UAS-Tkv^ACT^* flies. Error bars represent Mean ± SEM, *p <0.05; **p <0.01; ***p <0.001; ****p <0.0001 (two-way ANOVA). (X-Y) Midguts stained with anti-AstC (blue), anti-NPF (green), anti-CCHa2 (red) and anti-Tk (white) in the following genotypes of flies driven by *esg-Gal4^ts^* for 10 days since 48h APF: Control (X-X’) and *UAS-Tkv^ACT^* (Y-Y’). Scale bar in (A): 300 μm; (C): 100 μm; others: 50 μm.

We inhibited BMP signaling activity by overexpressing *Dad*, a BMP pathway negative regulator, or RNAi of BMP pathway effector Mad (*Mad-IR*) in ISCs started since early pupae stage (48h APF). *Dad-LacZ* expression was downregulated in the R1, R3 and R4a regions (**Figure 5—figure supplement 2D-E**). Consistent with previous findings on the role for BMP signaling in copper cell specification ^33^, a significant reduction of 2B10^+^ (monoclonal antibody against Cut) Copper cell number were observed in the R3 region following BMP inhibition (**Figure 5B-C**). We found that the proportion of CCHa2^+^ class III EEs in R2 region were increased (**Figure 5T**), and even were found to ectopically appeared in R1a and R3, where they should be absent (**Figure 5F-H, 5J-L, 5S, 5U and Figure 5—figure supplement 2A-B**). The proportion of these ectopic class III EEs was similar to that in the R2 region (**Figure 5S-U**). These results suggest that inhibition of BMP signaling partially induced pupal ISCs in R1a and R3 to adopt a cellular identity characteristic of R1b-R2 regions, thereby influencing EE subtype formation.

We further overexpressed an active form of BMP signaling effector Tkv (*Tkv^ACT^*) in pupal ISCs and observed upregulation of *Dad-LacZ* reporter expression throughout the entire midgut. Moreover, the cooper cell maker 2B10 appeared sporadically in the epithelial cells in R4 (**Figure 5D-E**), and the EE subtype composition was significantly altered in all midgut compartments. Normally, CCHa2 expression can be found in class III EEs in the R2 region (**Figure 5M**) and class I EEs in the R4b region (**Figure 5N**). However, in *UAS-Tkv^ACT^* guts, CCHa2 expression was virtually absent in all regions (**Figure 5I, 5P-Q, 5S-V and Figure 5—figure supplement 2C**) except R4c, where CCHa2^+^ but TK^-^ and AstC^-^ EEs, characteristics of class III EEs, appeared (**Figure 5O, 5R and 5W**). In all regions except R4c, EEs were all changed to a 1:1 composition of class I and class II EEs (**Figure 5S-W and Figure 5—figure supplement 2G-I**). Moreover, NPF, which is normally expressed in class II EEs in R2-R3, was found to be expressed in all Tk^+^ class II EEs at R4b region in these BMP overactivated midgut (**Figure 5X-Y**), while peptide hormones specifically expressed in EEs at the posterior midgut, including DH31 (**Figure 5—figure supplement 2J-M**) and AstA (**Figure 5—figure supplement 2N-Q**), were not ectopically expressed at R2 and R4b regions, indicating that EEs in these regions have assumed the identity from R3, and the loss of CCHa2 expression precludes their transformation into the identity from R2. Collectively, these results suggest that BMP signaling negatively regulates the specification of CCHa2^+^ EEs in most gut regions and plays an essential role in the establishment of regional EE identity and diversity.

### Manipulating WNT or BMP pathway activity in adult ISCs has limited effect on regional EE diversity

So far, we have demonstrated that WNT and BMP signaling could modulate regional pupal ISC identity and thereby facilitating EE subtype specification in different gut regions. Previous studies have shown that regional ISC identity was established during a defined time window of metamorphosis. Regulating BMP signaling during the early pupal stage can transform ISC and enterocyte identity, while this transformation cannot be induced in adult midgut ^34^. To further investigate whether regional EE diversity could be changed in adult flies, we performed WNT and BMP signaling activity manipulation in adult ISCs by using *esg-Gal4^ts^*. We transferred 5-day-old adult flies to 29℃ for 7 days, and then fed them with 5% DSS sucrose solution for 3 days and transferred back to standard food for 1-2 days, in order to induce stress and promote ISC proliferation to generate new EEs.

We observed that inhibiting either WNT signaling or BMP signaling did not alter the EE subtype composition in R1a (**Figure 6—figure supplement 1A, F and K**). However, inhibiting BMP signaling could sporadically induce a few class III EEs in R3 (**Figure 6—figure supplement 1B and G**). On the other hand, activating WNT signaling could induce a small portion of Tk^+^ class II EEs in R4b (**Figure 6—figure supplement 1D and L**). Activating BMP signaling did not significantly influence the EE subtype composition in R2 and R4b, but could induce a very small portion of class III EEs in the R4c region (**Figure 6—figure supplement 1C-E and H-J**). These subtle alterations in the composition of EE subtypes suggest that, alterations in WNT and/or BMP signaling activity within adult ISCs have a limited impact on the regional identity of ISCs responsible for generating EE subtypes.

### Manipulating WNT or BMP pathway activity in the differentiated EEs has a mild effect on their subtype identity

We have previously shown that the differentiated EEs in adult midgut remain to be plastic as changes of their TF code can switch one EE subtype into another ^49^. To test if the regional EE identity can be altered by manipulating BMP or WNT signaling activities within differentiated EEs, we inhibited or activated pathway activities specifically in EEs by using *prosV1-Gal4*.

We observed that the inhibition of BMP signaling led to ectopic appearance of a small portion of class III EEs in R3, but not in other regions (**Figure 6A-B, 6F-G and 6M-T**). The activation of BMP signaling led to the loss of class III EEs in R2 (**Figure 6C, 6H and 6N-O**) and the ectopic appearance of class III EEs in R4c (**Figure 6E, 6J and 6S**). However, CCHa2^+^ class I EEs distributed in the posterior midgut were not affected (**Figure 6D and 6I**). Basically, the activation of BMP signaling caused a tendency of transforming into the composition pattern of class I/II EE throughout the whole gut regions except R4c (**Figure 6M-T**).

**Figure 6.**
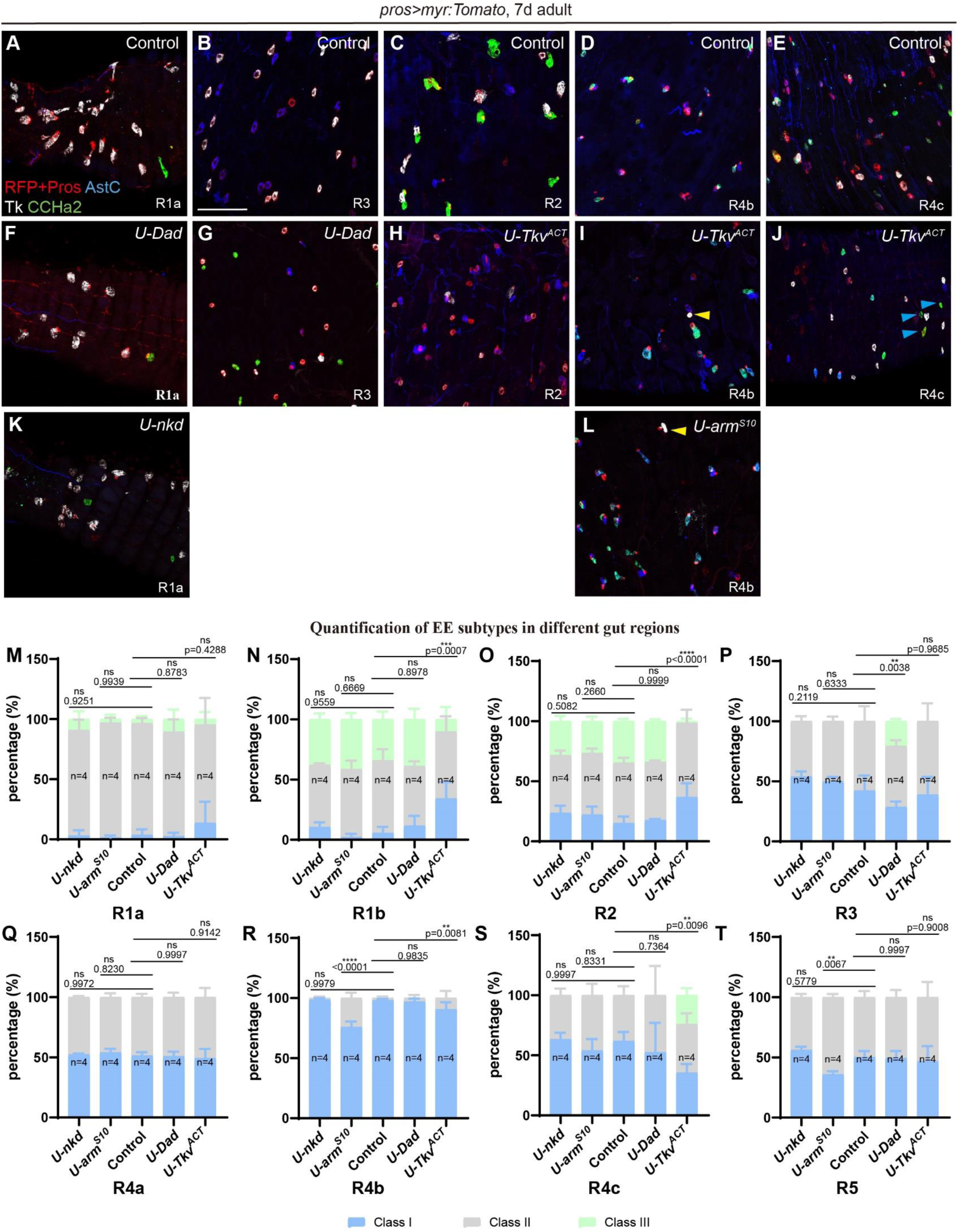
Manipulating WNT and BMP signaling in differentiated EEs has a mild impact on the regional specification of class II and III EEs. (A-L) Midguts stained with anti-Pros (red), anti-AstC (blue), anti-Tk (white) and anti-CCHa2 (green) in the following genotypes of flies driven by *pros-Gal4*: Control (A-E), *UAS-Dad* (F-G), *UAS-Tkv^ACT^* (H-J), *U-nkd* (K) and *U-arm^S^*^10^ (L). Yellow arrow heads in (I and L) represent ectopic class II EEs in R4b region, and blue arrow heads in (J) represent ectopic class III EEs in R4c region. Scale bar represents 25 μm. (M-T) Quantification of the three major EE subtype percentage in different gut regions of *UAS-nkd*, *UAS-arm^S^*^10^, Control, *UAS-Dad and UAS-Tkv^ACT^* flies. Error bars represent Mean ± SEM, **p <0.01; ***p <0.001; ****p <0.0001 (two-way ANOVA).

As for WNT signaling, inhibition of WNT activity did not significantly influence the EE composition in the whole gut (**Figure 6A, 6K and 6M-T**), but activation of WNT signaling induced ectopic appearance of a small portion of Tk^+^ class II EEs in R4b, and increased the percentage of class II EEs in R5 (**Figure 6D, 6L and 6M-T**). These phenotypes are similar but slightly weaker to those generated in ISCs initiated from the pupal stage. Therefore, BMP and WNT signaling gradients are not only important for establishing regional ISC identity and consequently contributing to the regional EE subtype diversity, but also play a role in maintaining regional diversity of differentiated EEs in adult midgut, reflecting the high plasticity of EEs.

### A combinatorial activity of WNT and BMP signaling gradients orchestrates regional EE diversity

Since both WNT and BMP signaling pathways contribute to the diversity of pupal ISCs and regional EE subtypes, we investigated whether specific combinations of WNT and BMP activity levels play a key role in shaping regional ISC identity and EE diversity. We aligned and compared the distribution of EE subtypes and EEP division modes with the activation level of WNT and BMP signaling pathways in each region along the entire midgut (**Figure 7M**). We found that in R3-R4a and R4c-R5, where WNT and BMP signaling activity are relatively high, EEP divides asymmetrically to generate class II/I EE subtypes. Inhibition of BMP signaling caused the ectopic appearance of class III EEs in R3, transforming the EE subtype composition into that of the R2 region (**Figure 5J-L, 5U and Figure 5—figure supplement 2D-E**), and activation of BMP signaling widely induced the transformation of EE subtype composition into class II/I EE subtypes in each compartment (**Figure 5S-W and Figure 5—figure supplement 2F-I**), indicating that a high level of BMP signaling induces EEP to divide asymmetrically and generate the composition of class II/I EE subtypes.

To further test this hypothesis, we simultaneously induced the activation or inhibition of WNT and BMP signaling in pupal ISCs, and the induction time still started since early pupal stage (48h APF). We found when the BMP signaling was activated, regardless of whether combined with the activated or inhibited WNT signaling, the resulting phenotypes were similar to observed upon activating BMP signaling alone (**Figure 5I, 5P-R, 5S-W, 7A-F, S9C and Figure 5—figure supplement 2F-I**), a condition under which a broad induction of class II/I EE subtype compositions occurred. Conversely, when the BMP signaling was inhibited, regardless of whether it was combined with the activated or inhibited WNT signaling, the phenotypic changes in the R3 region were similar to those of inhibiting BMP signaling alone (**Figure 5J-L, 5U, 7G-H and 7J**), a condition under which all three EE subtypes ectopically appeared in R3. These results suggest that, a high level of BMP signaling favors to induce asymmetric cell division of EEP to generate the composition of class II/I EE subtypes (**Figure 7N**).

**Figure 7.**
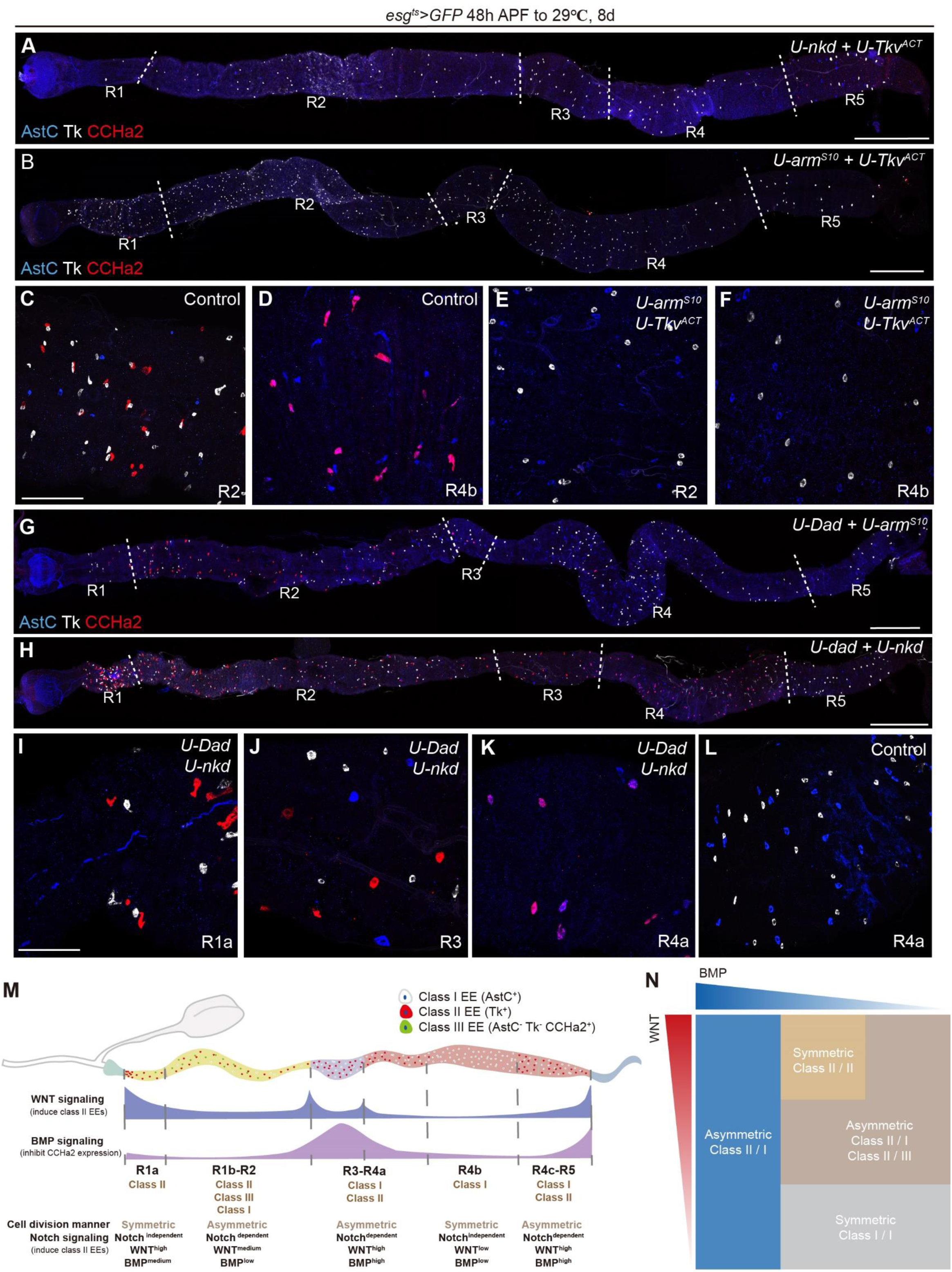
WNT and BMP signaling cooperatively regulate regional EE diversity. (A) Midguts stained with anti-AstC (blue), anti-Tk (white) and anti-CCHa2 (red) in *esg-Gal4^ts^>U-nkd, U-Tkv^ACT^* flies shift to 29 ℃ for 8 days since 48h APF. (B-F) Midguts stained with anti-AstC (blue), anti-Tk (white) and anti-CCHa2 (red) in the following genotypes of flies driven by *esg-Gal4^ts^* for 8 days since 48h APF: Control (C-D) and *UAS-arm^S^*^10^*, UAS-Tkv^ACT^* (B and E-F). (G) Midguts stained with anti-AstC (blue), anti-Tk (white) and anti-CCHa2 (red) in *esg-Gal4^ts^>U-Dad, U-arm^S^*^10^ flies shift to 29 ℃ for 8 days since 48h APF. (H-L) Midguts stained with anti-AstC (blue), anti-Tk (white) and anti-CCHa2 (red) in the following genotypes of flies driven by *esg-Gal4^ts^* for 8 days since 48h APF: Control (L) and *UAS-Dad, UAS-nkd* (H-K). (M) A schematic diagram describing the function of WNT and BMP gradients in regulating EEP division modes. (N) A schematic diagram on the relationships among EE subtype distribution pattern, WNT and BMP signaling activity and the EEP division modes along the fly midgut. Scale bar in (C) and (I): 50 μm; others: 300 μm.

We noticed that the composition of class II/I EE subtypes exist in the R3-R4a and R4c-R5 regions, but their peptide hormone profiles are significantly different ^12^. Inhibiting BMP signaling can transform the characteristics of R3 region, while activating BMP signaling widely induces other gut compartments to acquire the characteristics of the R3 region (**Figure 5S-W**), suggesting that a high level of BMP signaling establish the characteristics of the R3 region, while the characteristics of the R5 region may require precise activation levels of WNT and BMP signaling, or the involvement of additional unidentified mechanisms.

Next, we focused on the R1a region, where the WNT signaling is high, while BMP signaling is moderate or low, and EEP undergoes symmetric cell division to generate class II EEs (**Figure 7M**). We have found that inhibiting WNT signaling could induce the ectopic appearance of class III EEs in R1a (**Figure 4E-G and Figure 4—figure supplement 2F-G**), leading to asymmetric cell division of EEP to produce class II/III EE subtypes, which indicated that high level of WNT signaling determined the characteristic of R1a region. However, inhibition of BMP signaling also results in a similar phenotype in R1a (**Figure 5G-H and Figure 5—figure supplement 2D-E**), suggesting the involvement of BMP signaling. Indeed, inhibiting both WNT and BMP signaling simultaneously also led the ectopic appearance of class III EEs in R1a (**Figure 7H-I**). However, through genetic manipulation, we were unable to induce EEs in other gut regions to acquire the characteristics of R1a, perhaps because the establishment of the characteristic of R1a might require a precise combination of WNT and BMP activities or other signaling pathways, that promote the generation of class II/II EEs via symmetric division of EEPs (**Figure 7N**).

Finally, we concentrated on the R4b region, in which the activities of both WNT and BMP signaling are low, and EEP undergoes symmetric division to generate class I/I EEs (**Figure 7M**). We have found that activating WNT signaling caused composition of EE subtypes R4b to transform into the composition pattern of class II/I subtypes (**Figure 4D, 4I and Figure 4—figure supplement 2H**), and similar in BMP activated midguts (**Figure 5Q. 5V and Figure 5—figure supplement 2F**). To further test the synergistic effect of WNT and BMP signaling, we simultaneously inhibited WNT and BMP signaling in pupal ISCs, and found that CCHa2^+^ class I EEs ectopically appeared in the R4a region, and class II EEs were completely lost, resulting in a pattern that all EEs became class I EEs (**Figure 7K-L**). Based on these observations, we propose that the combination of low WNT and BMP signaling level favors the generation of class I/I EEs from symmetric division of EEPs, a pattern normally found in the R4b region (**Figure 7N**).

To further elucidate how Notch signaling coordinates regional EE identity with WNT and BMP signaling, we genetically manipulated WNT and BMP activities in the context of Notch loss-of-function (**Figure 7—figure supplement 1**). Notch depletion triggered rapid expansion of gut epithelial cells during late pupal and early adult stage, accompanied by the generation of large numbers of EEs, many of which remained immature, particularly in the R5 region (**Figure 7— figure supplement 1**). Further suppression of WNT activity (**Figure 7—figure supplement 1C and M**) or enhancement of BMP activity (**Figure 7—figure supplement 1B-D and Q-S**) exacerbated gut dysplasia and tumorigenesis (**Figure 7—figure supplement 1B, G and V**). Notably, Notch depletion alone did not alter the proportions of class I or class III EEs in R2, nor did it affect the proportion of CCHa2^-^ or CCHa2^+^ class I EEs in R4 (**Figure 3—figure supplement 2**). However, combined genetic manipulation of Notch and BMP signaling revealed that elevated BMP activity significantly suppressed CCHa2 expression in both R2 and R4 (**Figure 7—figure supplement 1A-D and P-S**), whereas BMP inhibition enhanced CCHa2 expression (**Figure 7—figure supplement 1A-D and U-X**). Collectively, these findings indicate that BMP signaling represses CCHa2 expression and may additionally modulate asymmetric cell division in the R2 and R4 regions.

Collectively, we propose that by default, class II, and I or III can be equally generated from ISCs via EEPs though Notch dependent (class II) and Notch-independent (class I and III) asymmetric cell divisions. WNT signaling activity favors the generation of class II EEs, while BMP signaling activity favors the generation of class I and against the generation of class III EEs (**Figure 7M**). As a result, the manifestation of EE subtype pattern in a specific gut region is, at least partially, governed by the intercalating levels of WNT and BMP signaling gradient emanating from the compartment boundaries (**Figure 7N**).

## Discussion

Our study suggests a model for the establishment of regional EE diversity in the *Drosophila* midgut: The WNT and BMP morphogen gradients, which emanated from compartment boundaries during metamorphosis and adulthood, act globally and locally (interacting with Notch) to regulate the regional patterning of EE subtypes (**Figure 7M-N**). We propose that a combined activity of WNT and BMP gradients at specific intercalated sites determines the cell division mode of EEPs and consequently influence the cell fate of EE progenies: high BMP activity (BMP^high^) promotes class I and II EE specification through Notch-dependent asymmetric EEP division (R3); high WNT activity combined with moderate BMP signaling (WNT^high^ and BMP^medium^) leads to the specification of class II EEs via symmetric EEP division (R1a); moderate levels of both WNT and BMP signaling (WNT^medium^ and BMP^medium^) facilitate the generation of all three EE subtypes through two distinct Notch-dependent asymmetric EEP divisions (R1b-R2); and low levels of both pathways (WNT^low^ and BMP^low^) favor the specification of class I EEs through symmetric EEP division (R4b).

However, several observations cannot be fully explained by the relative activities of WNT and BMP signaling. For example, the distinct expression patterns of CCHa2 and AstC in regions R1b–R2 and R4b–R4c, as well as the ectopic expression of CCHa2 in the R4c region under BMP hyperactivity, suggest additional regulatory influences. These findings indicate that other signaling pathways may be involved in specifying regional EE identity, particularly in distinguishing anterior from posterior regions. It is also worth noting that intrinsic mechanisms—such as region-specific transcription factor codes for EEs ^12,41^—may not be fully overridden by forced activation or inhibition of WNT and/or BMP signaling using the approaches employed in this study, potentially contributing to incomplete penetrance.

Our findings from the spatiotemporal manipulation of WNT and BMP signaling pathway further imply the significance of the intrinsic mechanisms—specifically, regional identity of ISCs—in determining regional EE diversity. This regional identity of ISCs is established during metamorphosis and cannot be efficiently altered after eclosion by manipulating WNT and BMP signaling activities. Additionally, ISCs located in different regions of the midgut surprisingly share an almost identical transcriptomic profile ^50^. These observations suggest that an epigenetic mechanism may underlie both the establishment and maintenance of regional ISC identity, as well as the regional characteristics of differentiated cells in adulthood. These results are consistent with previous findings indicating that BMP signaling overactivation enables efficient acquisition of gastric stem cell identity in the R3 region at the early pupal stage, but not in adult stages ^34^. In cases of injury or aging stress, extrinsic signals of the midgut undergo significant changes, such as the widespread activation of BMP signaling induced by bleomycin or pathogenic bacteria challenge ^47,51^. If the regional identity of ISCs is too readily and rapidly modifiable, it is conceivable that this could lead to disorders of intestinal epithelial compartmentation and potentially severe consequences such as metaplasia.

Interestingly, we found that manipulating WNT and BMP signaling in differentiated EEs can modify their regional identities to a certain extent. Our previous work has demonstrated that EEs exhibit a notable degree of cellular plasticity: alterations in their TF code—sometimes by modifying the expression of a single TF—can effectively switch their subtype identity ^41^. Extrinsic signaling cues, such as WNT and BMP gradients, may act to fine-tune these TF networks, thereby influencing EE identity. Thus, to a certain extent, differentiated EEs appear responsive to WNT and BMP morphogens, which may not only contribute to the maintenance of regional subtype identity under homeostatic conditions but also facilitate adaptive changes in cellular identity and peptide hormone expression in response to environmental and physiological demands.

Apart from the A/P axis, the mammalian gastrointestinal tract is more complex than that of *Drosophila*, with the addition of the axis along the villus and crypt. However, the conservation of intrinsic and extrinsic factors regulating ISC differentiation and EE subtype specification across *Drosophila* and mammals ^6^ further suggest that, WNT and BMP signaling may represent conserved mechanism for regulating regional EE diversity in both organisms. Studies have shown that EEs originating from the crypt can switch their subtype identity as they migrate upward along the villus-crypt axis, a phenomenon attributed to the BMP signaling gradients ^52^. Furthermore, treatment with BMP agonists in human intestinal organoids has been found to promote the generation of EE subtypes typically localized to the upper villus region, where BMP signal is the highest ^52^. Interestingly, despite the apparent plasticity of EEs, intestinal cells within cultured organoids or those transplanted into the colon, retain their original regional identity even after prolonged culture or transplantation ^9,53–55^. Collectively, these observations support the notion that the regional ISC identity is maintained postnatally through intrinsic mechanisms, while EEs remain plastic and responsive to extrinsic morphogen signals in both *Drosophila* and mammals. Our recent scRNA-seq analysis of *Drosophila* EEs has further classified the three major EE classes into 10 distinct subtypes, each characterized by the expression of a unique combination of 2–4 peptide hormones^12^. This study establishes a foundation for enhancing the comprehension of the specification and regulatory mechanisms of EE subtypes in *Drosophila*. It is anticipated to contribute to the understanding of regional EE diversity and regulation in mammals and humans, thereby pave the way for the development of novel therapeutic strategies for enteroendocrine-related disorders and diseases, such as obesity and diabetes.

## Materials and Methods

### Fly strains and cultivation

The following fly strains were used in this study: *CCHa2-Gal4* (BDSC, #84602); *CCHa2-LexA* (BDSC, #84962); *Tk-Gal4* (BDSC, #84693); *AstC-Gal4* (BDSC, #84595); *GMR45D10-Gal4* (BDSC, #45323); *GMR50A12-Gal4* (BDSC, #47618); *GMR46B08-Gal4* (BDSC, #47361); *esg-GAL4,UAS-GFP* (gift from Shigeo Hayashi); *prosV1-Gal4* (gift from Bruce Edgar); *Wnt4-Gal4* (BDSC, #67449); *Fz3-RFP* (BDSC, #92358); *Dad-LacZ* (BDSC, #10305); *UAS-mCD8:RFP* (BDSC, #27398); *LexAOP-GFP, UAS-RFP* (BDSC, #32229); *LexAOP-mCherry* (BDSC, #52271); *UAS-Notch RNAi* (BDSC, #7078); *UAS-scute* (BDSC, #26687); *UAS-nkd.GFP* (BDSC, #25384); *UAS-Tcf^DN^* (BDSC, #4785); *UAS-Tcf RNAi* (BDSC, #26743); *UAS-arm^S^*^10^ (BDSC, #4782); *UAS-Dad.T* (BDSC, #98452); *UAS-Mad RNAi* (BDSC, #31315); *UAS-Tkv^ACT^* (gift from Chip Furguson); *Dl^Revf^*^10^ (BDSC, #6300); *Apc^Q^*^8^*, Apc2^G^*^10^ (gift from Mark Peifer); *82B FRT*; *hsflp, Act-Gal4,UAS-EGFP; 82B FRT-Gal80* were all obtained from BDSC. Fly stocks were cultivated on standard food with yeast paste added on the food surface and kept at 25 ℃ unless otherwise stated.

The Gal4/UAS/Gal80^ts^ or LexA-LexAOP systems were used to conduct conditional knocking down or overexpression in specific cell types ^56–58^. Unless otherwise stated, all crosses were performed at 18℃, and newly formed F1 pupae were collected and transferred to 29 ℃ to induce gene expression at 48h APF (corresponding to 25 ℃).

To induce gut stress and promote EE pair formation, liquid food containing 5% DSS (MP Biomedicals, #160110) and 5% sucrose (Xilong Scientific, #57-50-1) was added to extra thick filter paper sheet (Bio-Rad, #1703966) and supplied to adult flies for 2-3 days at corresponding temperature.

### Antibody generation

The mouse monoclonal anti-CCHa2, anti-DH31, anti-AstA and anti-NPF antibodies were custom made by Antibody Research Center in National Institute of Biological Sciences, Beijing. As described previously ^59–61^, a mature amidated CCHa2 peptide (GGGGSGCQAYGHVCYGGH-NH2), a full-length amidated DH31 peptide (CTVDFGLARGYSGTQEAKHRMGLAAANFAGGP-NH2), AstA peptide (NGGPGMKRLPVYNFGL-NH2) and full-length mature NPF (CSNSRPPRKNDVNTMADAYKFLQDLDTYYGDRARVRF-NH2) were synthesized and used for immunization separately. The peptides were injected into mice until the diluted raw sera showed immunostaining on *Drosophila* EEs. Following the fusion, hybridoma lines were screened by ELISA and immunostaining on *Drosophila* midgut, resulting in confirmed positives. The human monoclonal anti-CCHa2 was custom made by Biointron (Shanghai, China). variable region sequence of mAb was fished and recombinant expressed as human IgG.

### Immunostaining

Immunostaining of *Drosophila* midgut was performed as previously described ^45^. In brief, 10-15 adult female flies for each sample were dissected in PBS and then fixed in 4% paraformaldehyde for 30 min at room temperature, followed by dehydration in methanol (for 5 min) and rehydration in PBT solution (PBS containing 0.1% Triton X-100, 5 min each for 3 times). The primary antibodies were added into 300 μl 5% NGS-PBT solution and incubate with sample for overnight at 4 ℃. After washing three times using PBT, the secondary antibodies were added into 300 μl PBT and incubate with samples for 2 hours at room temperature, followed by staining of DAPI for 5 min. 80% glycerol was used to mount the samples and slides were kept in -20 ℃ freezer. Images were acquired on a Leica SP8 and Nikon AX inverted confocal microscopes. All images were assembled in Adobe Photoshop and Illustrator.

Primary antibodies used in this study were listed as follows: mouse anti-Pros (DSHB #MR1A; 1:300); rabbit anti-AstC ^29^ (gift from Dr. Jan A. Veenstra); guinea pig polyclonal anti-Tk ^62^ (gift from Dr. Eun Young Kim); rabbit polyclonal anti-β-galactosidase (Cappel, 0855976; 1: 6000); rabbit polyclonal anti-RFP (Rockland, 600-401-379; 1:400); rabbit monoclonal anti-GFP (Invitrogen, G10362; 1:600); mouse monoclonal anti-CCHa2 (this study; 1:300); human monoclonal anti-CCHa2 (this study; 1:300); mouse monoclonal anti-DH31 (this study; 1:300); mouse monoclonal anti-NPF (this study; 1:100); mouse monoclonal anti-AstA (this study; 1:200). Secondary antibodies used in this study include Alexa Fluor 405-conjugated donkey anti-mouse IgGs (Invitrogen, A148257; 1:300); Alexa Fluor 488-or 568-conjugated goat anti-mouse IgGs (Invitrogen, A11029, A11031; 1:300); Alexa Fluor 405-, 488-, 568- or Cy5-conjugated goat anti-rabbit IgGs (Invitrogen, A48254, A11034, A11036, A10523; 1:300); Alexa Fluor 488-, 568- or 647-conjugated goat anti-guinea pig IgGs (Invitrogen, A11073, A11075, A21450; 1:300); Alexa Fluor 633-conjugated goat anti-human IgGs (Invitrogen, A21091, 1:300) and DAPI (Sigma-Aldrich, 1μg/ml) was used for nuclei staining.

### Statistical Analysis

Cell number counting and mean fluorescent intensity were performed using Image J, and all quantifications were presented in the form of mean±SEM. GraphPad Prism 8 software was used to calculate p-values by unpaired student’s t-test or two-way Anova test.

## Acknowledgement

We thank Dr. Eun Young Kim for providing anti-Tk antibody, Dr. Jan-Adrianus Veenstra for anti-AstC antibody, Drs. Bruce Edgar, Chip Furguson, Shigeo Hayashi, Mark Peifer, Yi Rao for providing fly strains and the Bloomington *Drosophila* Stock Center (BDSC), the Tsinghua Fly Center, and Development Studies Hybridoma Bank (DSHB) and Antibody Center in NIBS for fly strains and antibodies, and members of the Xi laboratory for discussion and reading of the manuscript. This work was supported by National Key Research and Development Program of China (2020YFA0803502 and 2017YFA0103602 to R.X.) from the Chinese Ministry of Science and Technology. X.G. is supported by National Natural Science Foundation of China (Grant No. 32100595).

## Author Contributions

J.L., X.G. and R.X. designed the experiments. J.L. and X.G. performed the experiments. R.X. acquired funding. J.L., X.G. and R.X. wrote the paper.

---

**Figure 1—figure supplement 1.**
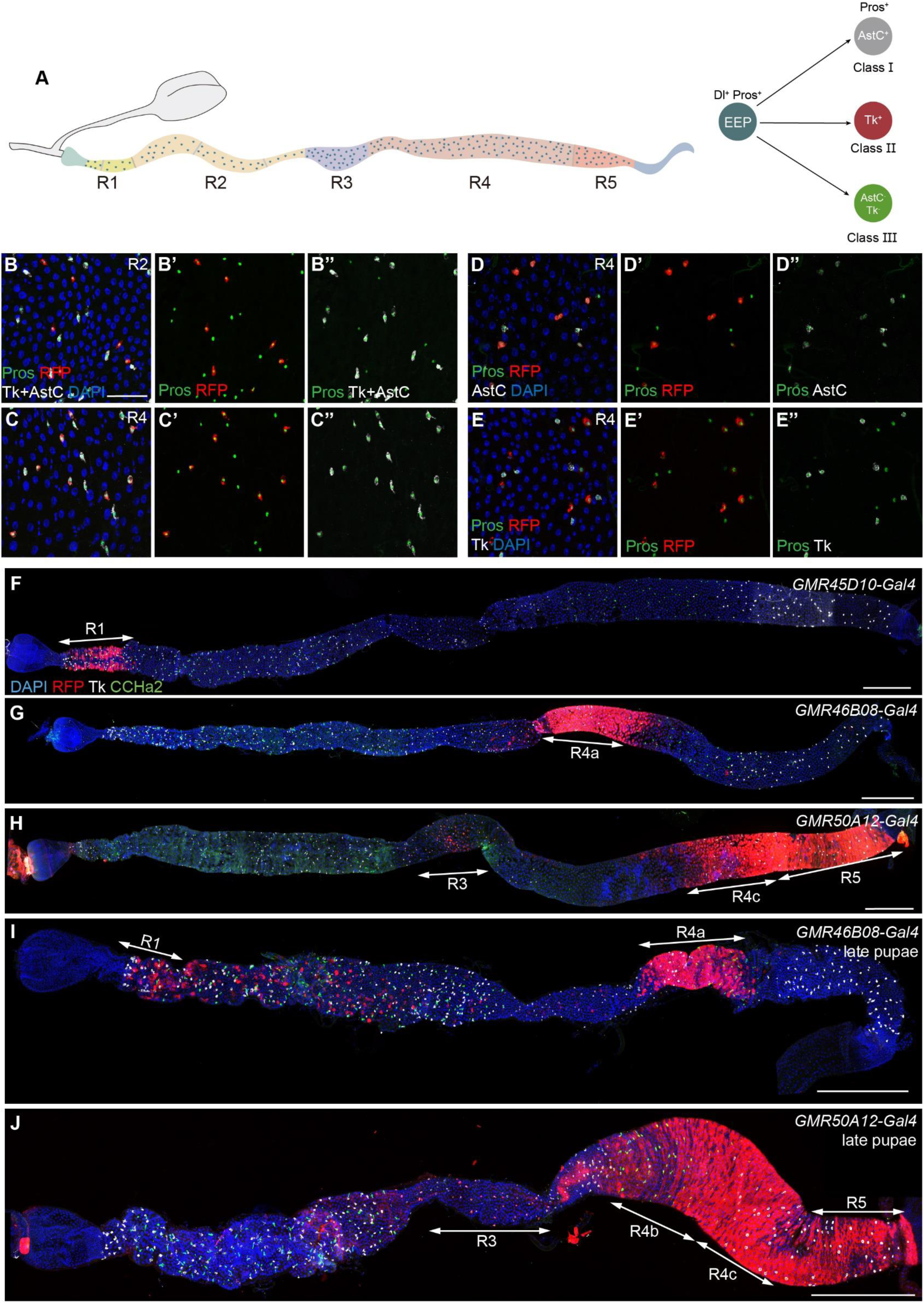
The spatial distribution of EE subtypes with highlighted compartmental boundaries in adult and pupal fly midgut. (A) A schematic diagram of *Drosophila* adult midgut compartmentation and three major EE subtypes. (B-C) Midguts of *CCHa2>RFP* flies stained with DAPI (blue), anti-Pros (green), anti-AstC and anti-Tk (white) antibodies. (D-D’’’) Midguts of *CCHa2>RFP* flies stained with DAPI (blue), anti-Pros (green) and anti-AstC (white) antibodies. CCHa2 colocalized with AstC in EEs in the R4 region. (E-E’’’) Midguts of *CCHa2>RFP* flies stained with DAPI (blue), anti-Pros (green) and anti-Tk (white) antibodies. (F) Midguts of *GMR45D10-Gal4>UAS-RFP* flies stained with DAPI (blue), anti-CCHa2 (green), anti-RFP (red) and anti-Tk (white) antibodies. (G) Midguts of *GMR46B08-Gal4>UAS-RFP* flies stained with DAPI (blue), anti-CCHa2 (green), anti-RFP (red) and anti-Tk (white) antibodies. (H) Midguts of *GMR50A12-Gal4>UAS-RFP* flies stained with DAPI (blue), anti-CCHa2 (green), anti-RFP (red) and anti-Tk (white) antibodies. (I) Late pupal midguts of *GMR46B08-Gal4>UAS-RFP* flies stained with DAPI (blue), anti-CCHa2 (green), anti-RFP (red) and anti-Tk (white) antibodies. (J) Late pupal midguts of *GMR50A12-Gal4>UAS-RFP* flies stained with DAPI (blue), anti-CCHa2 (green), anti-RFP (red) and anti-Tk (white) antibodies. Scale bar in (B): 25 μm; others: 300 μm.

**Figure 1—figure supplement 2.**
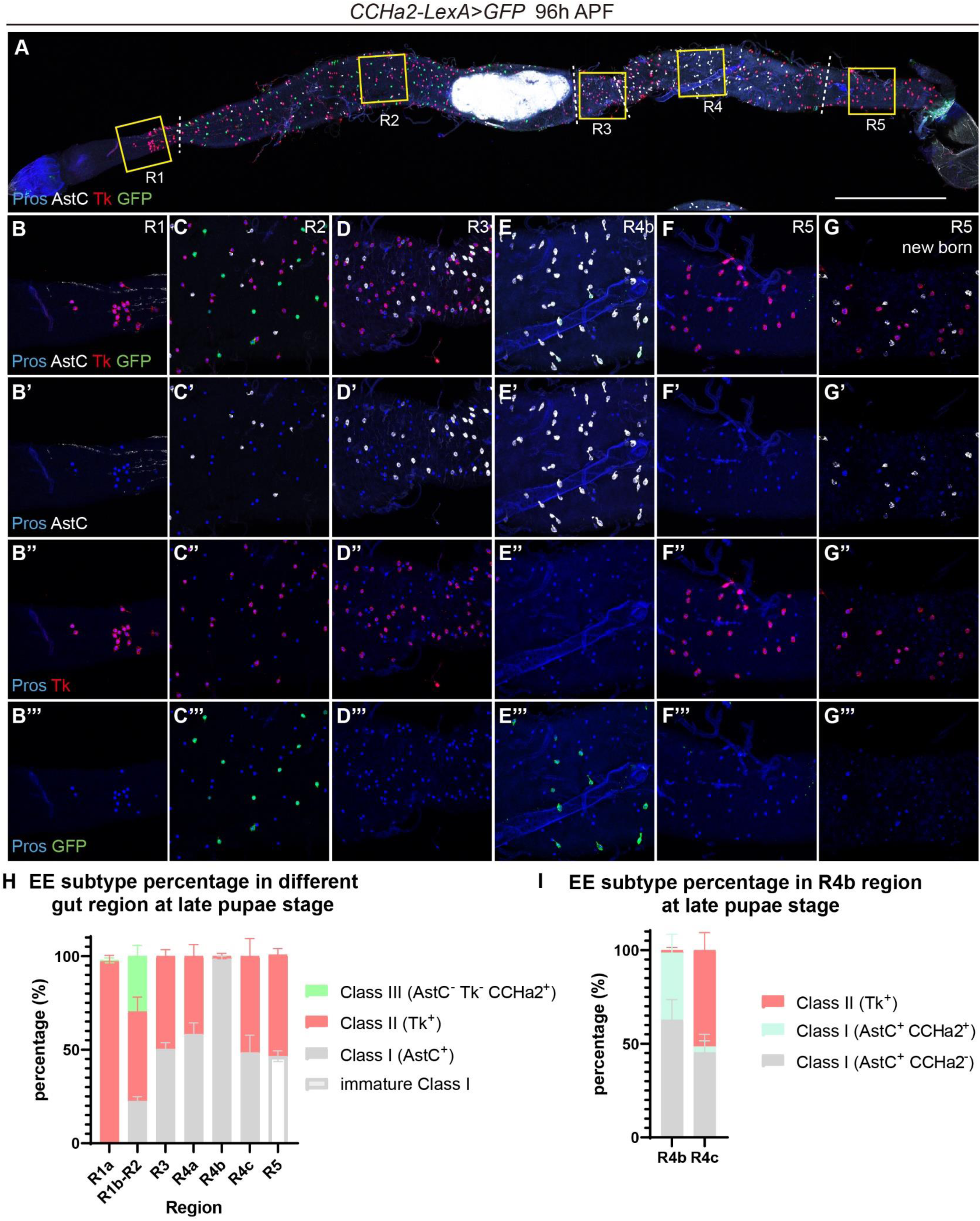
The distribution and composition of EE subtypes in different gut regions at late pupal stage. (A-F) Midguts of *CCHa2-LexA>GFP* flies stained with anti-Pros (blue), anti-AstC (white) and anti-Tk (red) antibodies before eclosion. Separated color channels for the insets from different gut regions in (A) were enlarged and displayed in (B-F), showing EE distribution characteristic of different gut regions. Scale bar represents 300 μm. (G) Midguts of *CCHa2-LexA>GFP* flies stained with anti-Pros (blue), anti-AstC (white) and anti-Tk (red) antibodies just after eclosion. (H) Quantification of the three major EE subtype percentage in different gut regions of flies before eclosion. Error bars represent Mean ± SEM, n=5. (I) Quantification of EE subtype percentage in the R4bc region of flies before eclosion. Error bars represent Mean ± SEM, n=5.

**Figure 2—figure supplement 1.**
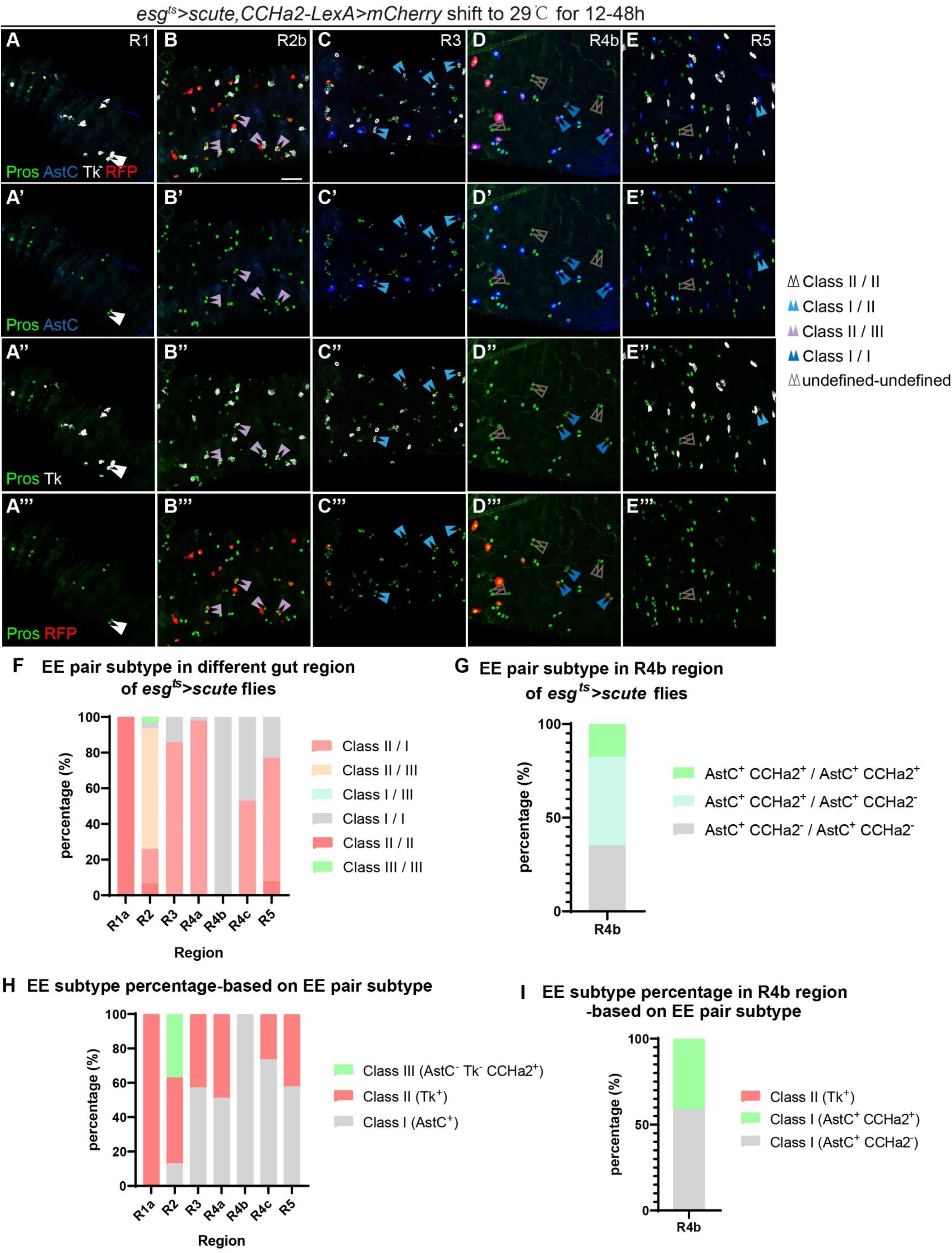
Characterization of *scute* overexpression-induced EE pairs in different gut regions. (A-E) Midguts stained with anti-Pros (green) and anti-AstC (blue) antibodies in *esg^ts^>scute, CCHa2-LexA>mCherry* flies that shift to 29 ℃ to induce expression for 12-48 hours. EE pairs were pointed by arrow head pairs. Scale bar represents 25 μm. (F) Quantification of different EE subtype pair percentage in different gut regions. (R1: n=4; R2: n=31; R3: n=21; R4b: n=17; R5: n=13) (G) Quantification of different EE subtype pair percentage in the R4b region. (n=64) (H) Speculation of EE subtype percentage in different gut regions by taking the percentage of EE subtype pairs from (F) to generate EEs. (I) Speculation of EE subtype percentage in the R4b region by taking the percentage of EE subtype pairs from (G) to generate EEs.

**Figure 3—figure supplement 1.**
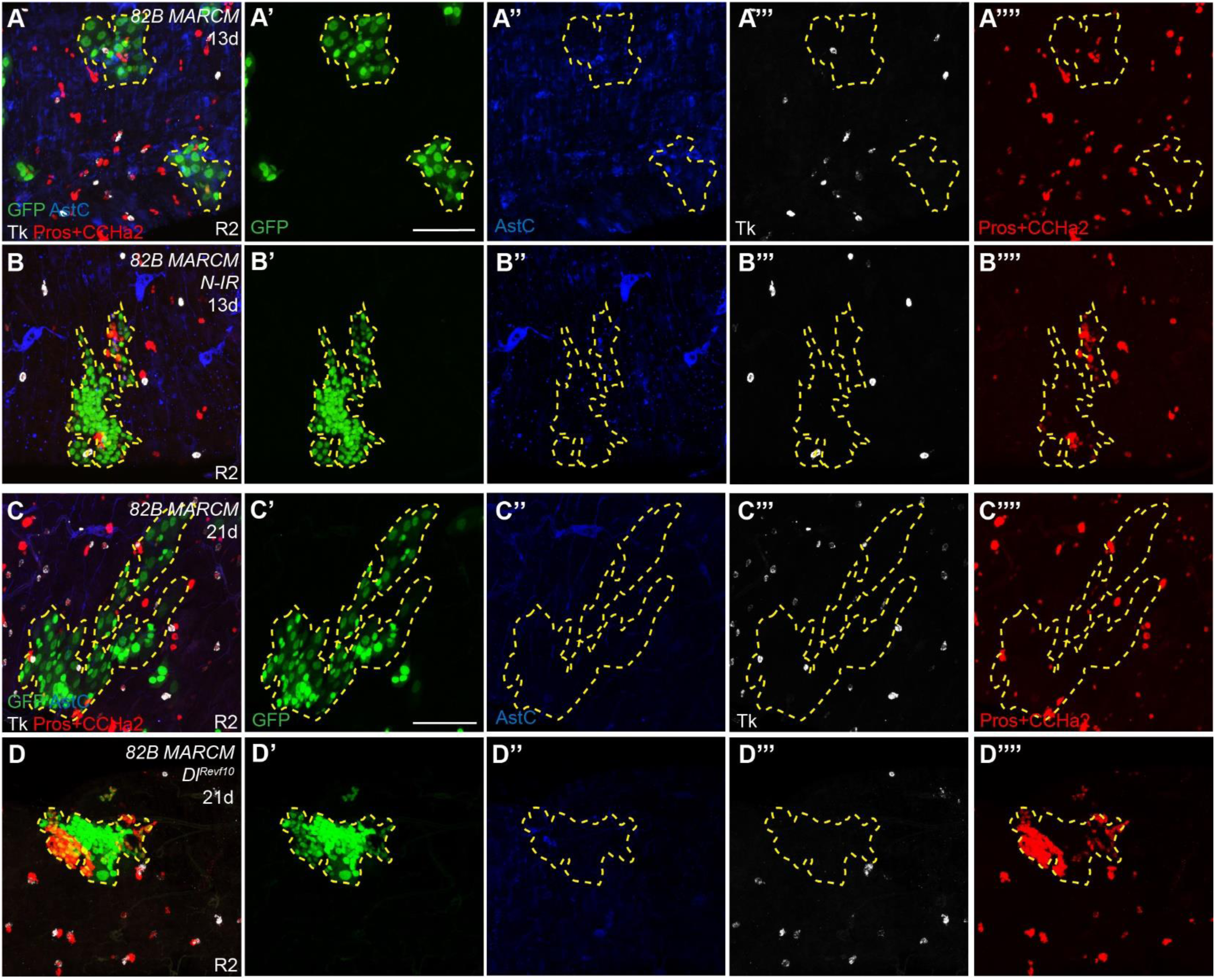
Notch is required for the specification of class II but not class I or III subtypes. (A-B) MARCM clones co-stained with anti-AstC (blue), anti-Tk (white) and anti-Pros, anti-CCHa2 (red) (Pros is expressed in EE nucleus and CCHa2 is expressed in cytoplasm, which can be distinguished by subcellular localization of the fluorescence signal). (A-A’’’) WT *FRT82B* clone, (B-B’’’) *N-IR FRT82B* MARCM. (C-D) MARCM clones co-stained with anti-AstC (blue), anti-Tk (white) and anti-Pros, anti-CCHa2 (red). (C-C’’’) WT *FRT82B* clone, (D-D’’’) *Dl^Revf^*^10^ *FRT82B* MARCM. Scale bars represent 25 μm.

**Figure 3—figure supplement 2.**
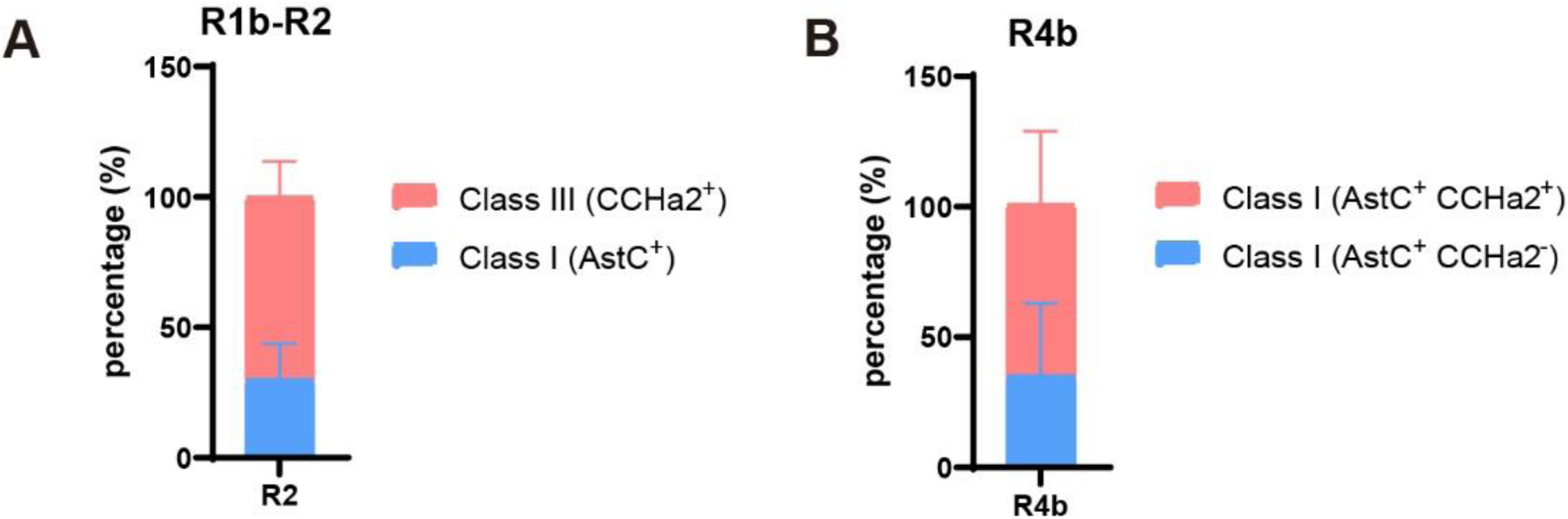
Loss of Notch biases EE cell fate specification. (A) Quantification of class I and class III EE subtype percentage in EE clusters from R1b-R2 region of *esg^ts^>N-IR* flies. Error bars represent Mean ± SEM, n=37. (B) Quantification of AstC^+^ CCHa2^-^ class I and AstC^+^ CCHa2^+^ class I EE subtype percentage in EE clusters from R4b region of *esg^ts^>N-IR* flies. Error bars represent Mean ± SEM, n=22.

**Figure 4—figure supplement 1.**
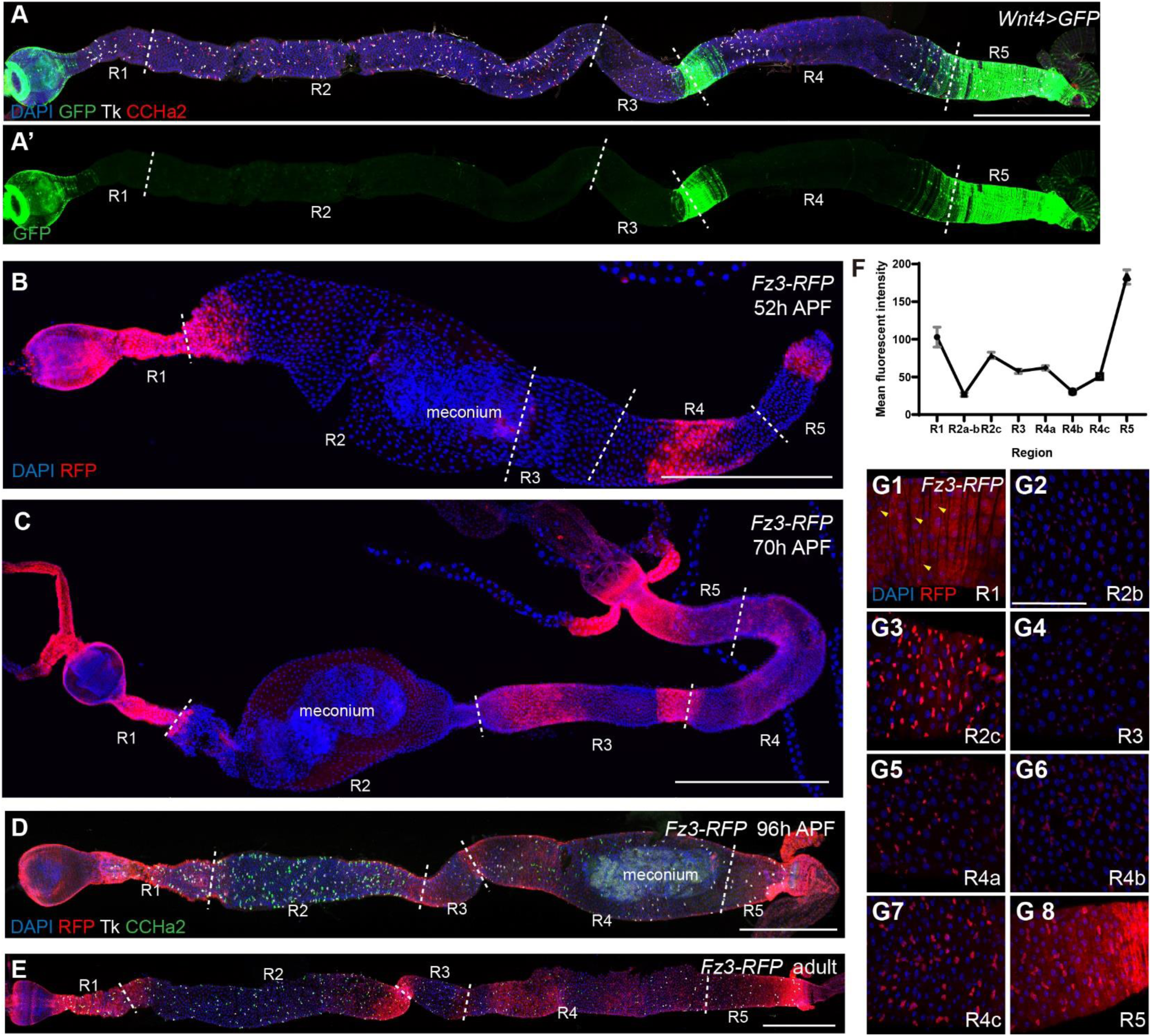
Characterization of WNT signaling pathway activity along the midgut. (A) Midguts of *Wnt4>GFP* flies stained with DAPI (blue), anti-GFP (green), anti-Tk (white) and anti-CCHa2 (red) antibodies after eclosion. (B-C) Midguts of *Fz3-RFP* flies stained with DAPI (blue) and anti-RFP (red) antibodies at 52h APF and 70h APF. (D-E) Midguts of *Fz3-RFP* flies stained with DAPI (blue), anti-RFP (red), anti-Tk (white) and anti-CCHa2 (green) antibodies at 96h APF (D) and 5 days after eclosion (E). (F) Mean fluorescent intensity of Fz3-RFP along the whole midgut at adult stage. Error bars represent Mean ± SEM, n=5. (G1-G8) Expression of *Fz3-RFP* in ISCs within different gut regions. Midguts of *Fz3-RFP* flies stained with DAPI (blue) and anti-RFP (red) antibodies 5 days after eclosion. ISCs in the R1 region were pointed by yellow arrow heads in (G1). Scale bar in (G): 50 μm; others: 300 μm.

**Figure 4—figure supplement 2.**
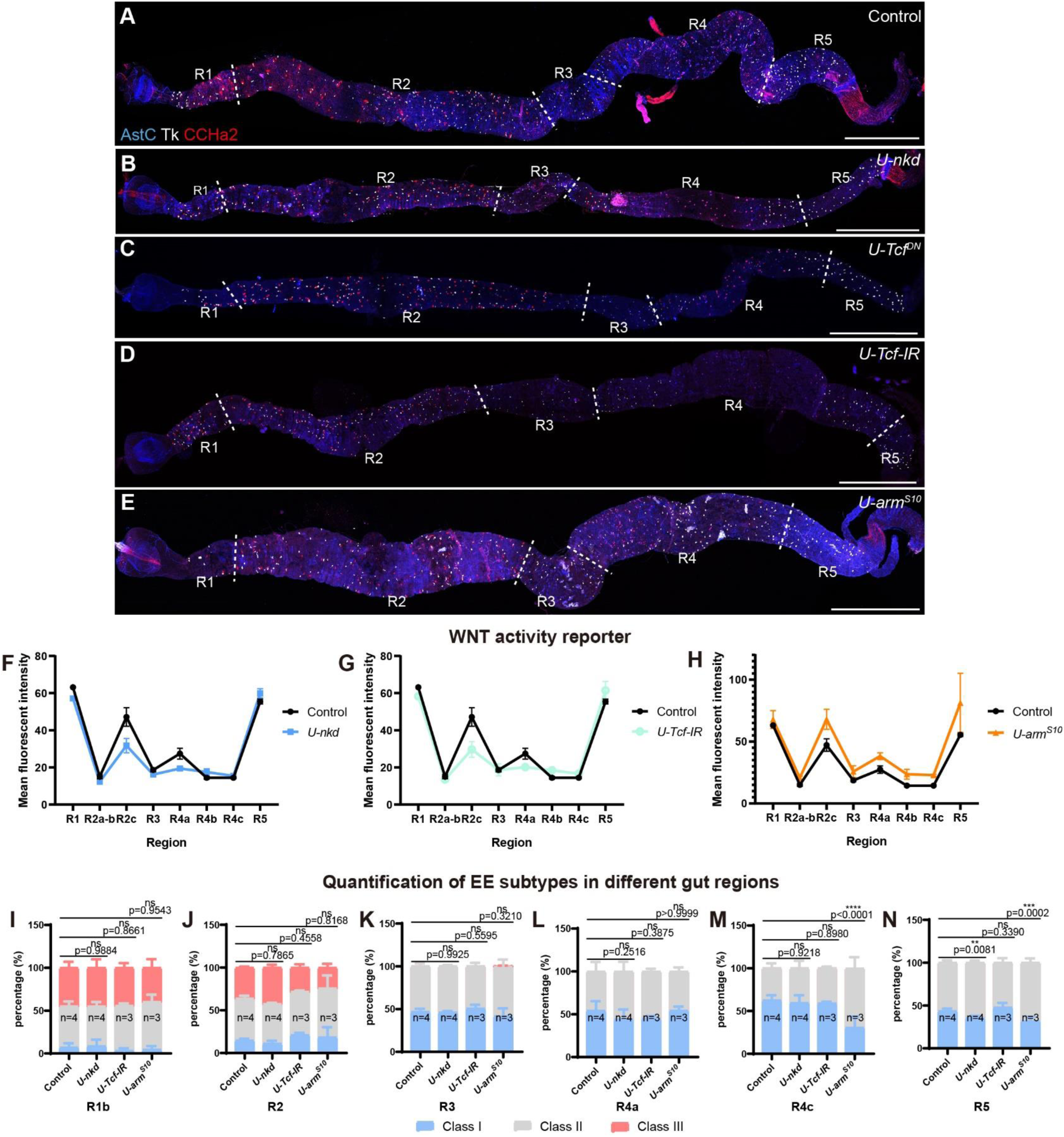
Impact of WNT signaling manipulation on regional EE subtype patterns. (A-E) Midguts stained with anti-AstC (blue), anti-Tk (white) and anti-CCHa2 (red) in the following genotypes of flies driven by *esg-Gal4^ts^* for 8 days since 48h APF: Control (A), *UAS-nkd* (B), *UAS-Tcf^DN^*(C), *UAS-Tcf RNAi* (D) and *UAS-arm^S^*^10^ (E). Scale bars represent 300 μm. (F-H) Mean fluorescent intensity of Fz3-RFP along the whole midgut at adult stage of Control, *UAS-nkd* (F), *UAS-Tcf RNAi* (G) and *UAS-arm^S^*^10^ (H) flies. Error bars represent Mean ± SEM, n=3. (I-N) Quantification of the three major EE subtype percentage in R1b (I), R2 (J), R3 (K), R4a (L), R4c (M) and R5 (N) of Control, *UAS-nkd*, *UAS-Tcf RNAi* and *UAS-arm^S^*^10^ flies. Error bars represent Mean ± SEM, **p <0.01; ***p <0.001; ****p <0.0001 (two-way ANOVA).

**Figure 5—figure supplement 1.**
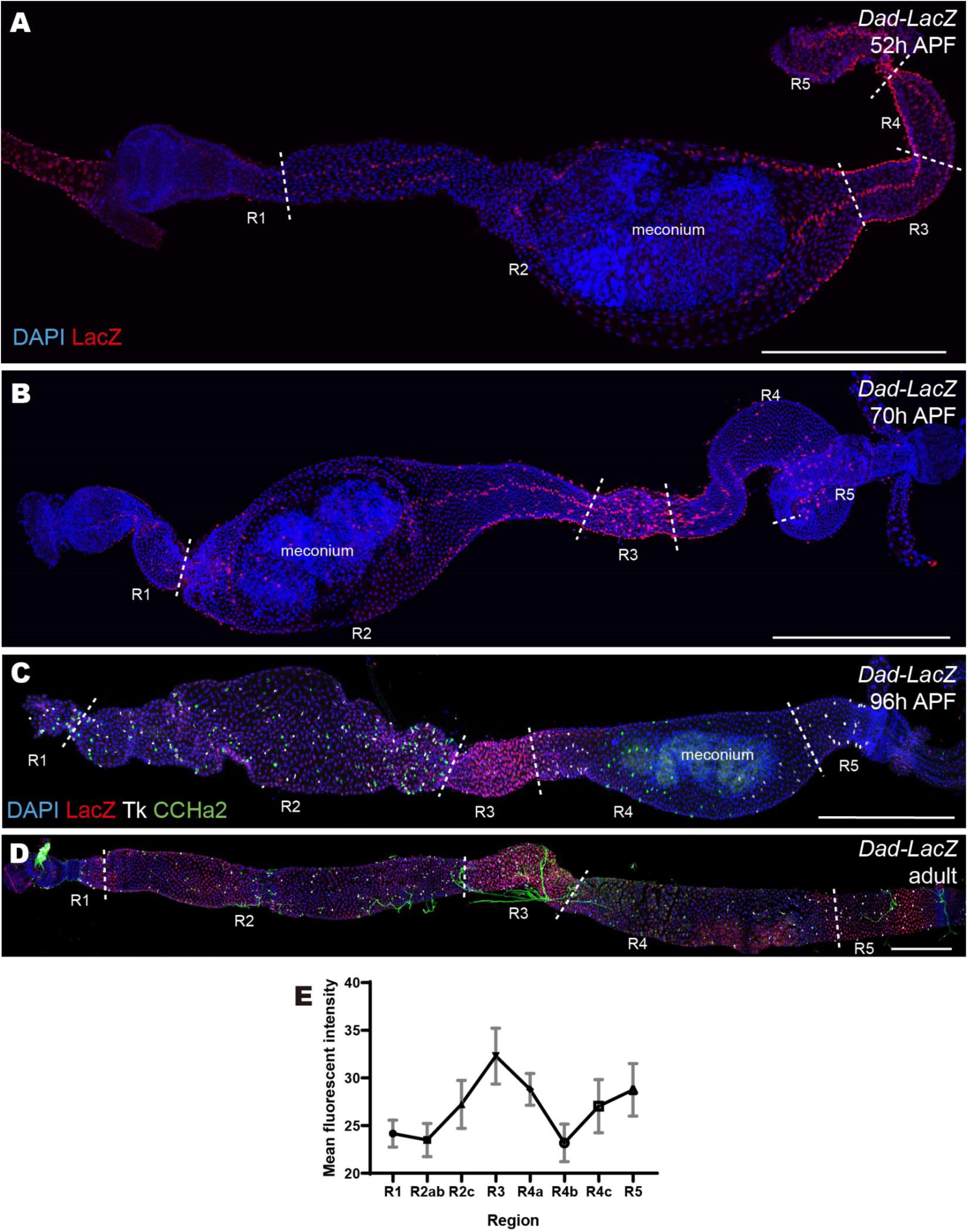
Characterization of BMP signaling pathway activity along the midgut. (A-B) Midguts of *Dad-LacZ* flies stained with DAPI (blue) and anti-LacZ (red) antibodies at 52h APF and 70h APF. (C-D) Midguts of *Dad-LacZ* flies stained with DAPI (blue), anti-LacZ (red), anti-Tk (white) and anti-CCHa2 (green) antibodies at 96h APF and 5 days after eclosion. (E) Mean fluorescent intensity of Dad-LacZ along the whole midgut at adult stage. Error bars represent Mean ± SEM, n=4. Scale bars represent 300 μm.

**Figure 5—figure supplement 2.**
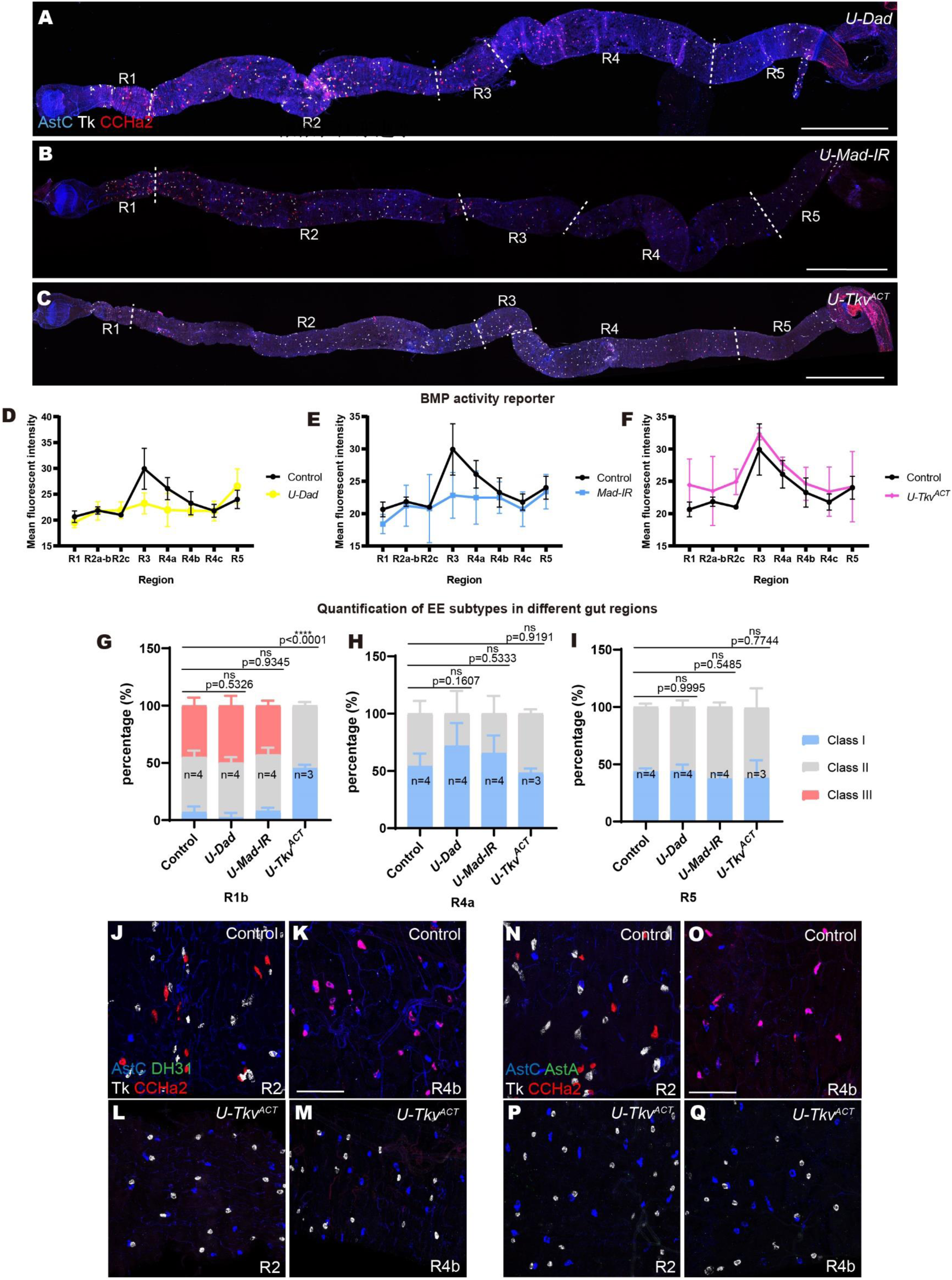
Impact of BMP signaling manipulation on regional EE subtype patterns. (A-C) Midguts stained with anti-AstC (blue), anti-Tk (white) and anti-CCHa2 (red) in the following genotypes of flies driven by *esg-Gal4^ts^* for 8 days since 48h APF: *UAS-Dad* (A), *UAS-Mad RNAi* (B) and *UAS-Tkv^ACT^* (C). (D-F) Mean fluorescent intensity of Dad-lacZ along the whole midgut at adult stage of Control, *UAS-Dad* (D), *UAS-Mad RNAi* (E) and *UAS-Tkv^ACT^* (F) flies. Error bars represent Mean ± SEM, n=3. (G-I) Quantification of the three major EE subtype percentage in R1b (G), R4a (H) and R5 (I) of *UAS-Dad*, *UAS-Mad RNAi* and *UAS-Tkv^ACT^* flies. Error bars represent Mean ± SEM, ****p <0.0001 (two-way ANOVA). (J-M) Midguts stained with anti-AstC (blue), anti-DH31 (green), anti-CCHa2 (red) and anti-Tk (white) in the following genotypes of flies driven by *esg-Gal4^ts^* for 10 days since 48h APF: Control (J-K) and *UAS-Tkv^ACT^* (L-M). (N-Q) Midguts stained with anti-AstC (blue), anti-AstA (green), anti-CCHa2 (red) and anti-Tk (white) in the following genotypes of flies driven by *esg-Gal4^ts^* for 10 days since 48h APF: Control (N-O) and *UAS-Tkv^ACT^* (P-Q). Scale bar in (A-C): 300 μm; (J-Q): 50 μm.

**Figure 6—figure supplement 1.**
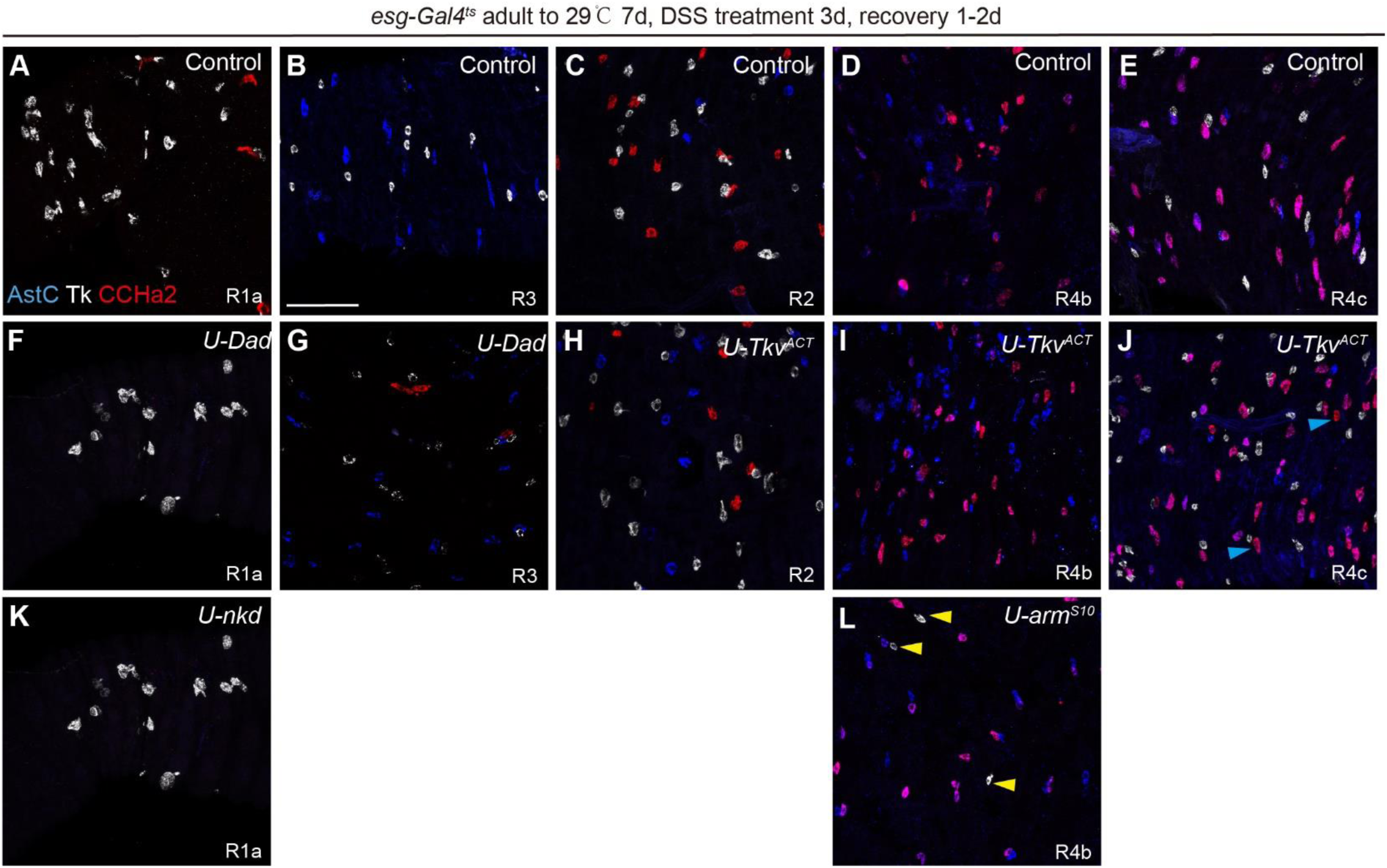
Manipulating WNT and BMP signaling in adult ISCs causes a mild effect on regional patterning of class II and III EEs. (A-L) Midguts stained with anti-AstC (blue), anti-Tk (white) and anti-CCHa2 (red) in the following genotypes of flies driven by *esg-Gal4^ts^* for 7 days and following treatment of 5% DSS for 3 days and recovery for 1 day at adult stage: Control (A-E), *UAS-Dad* (F-G), *UAS-Tkv^ACT^* (H-J), *U-nkd* (K) and *U-arm^S^*^10^ (L). Yellow arrow heads in (L) represent ectopic class II EEs in R4b region, and blue arrow heads in (J) represent ectopic class III EEs in R4c region. Scale bar represents 25 μm.

**Figure 7—figure supplement 1.**
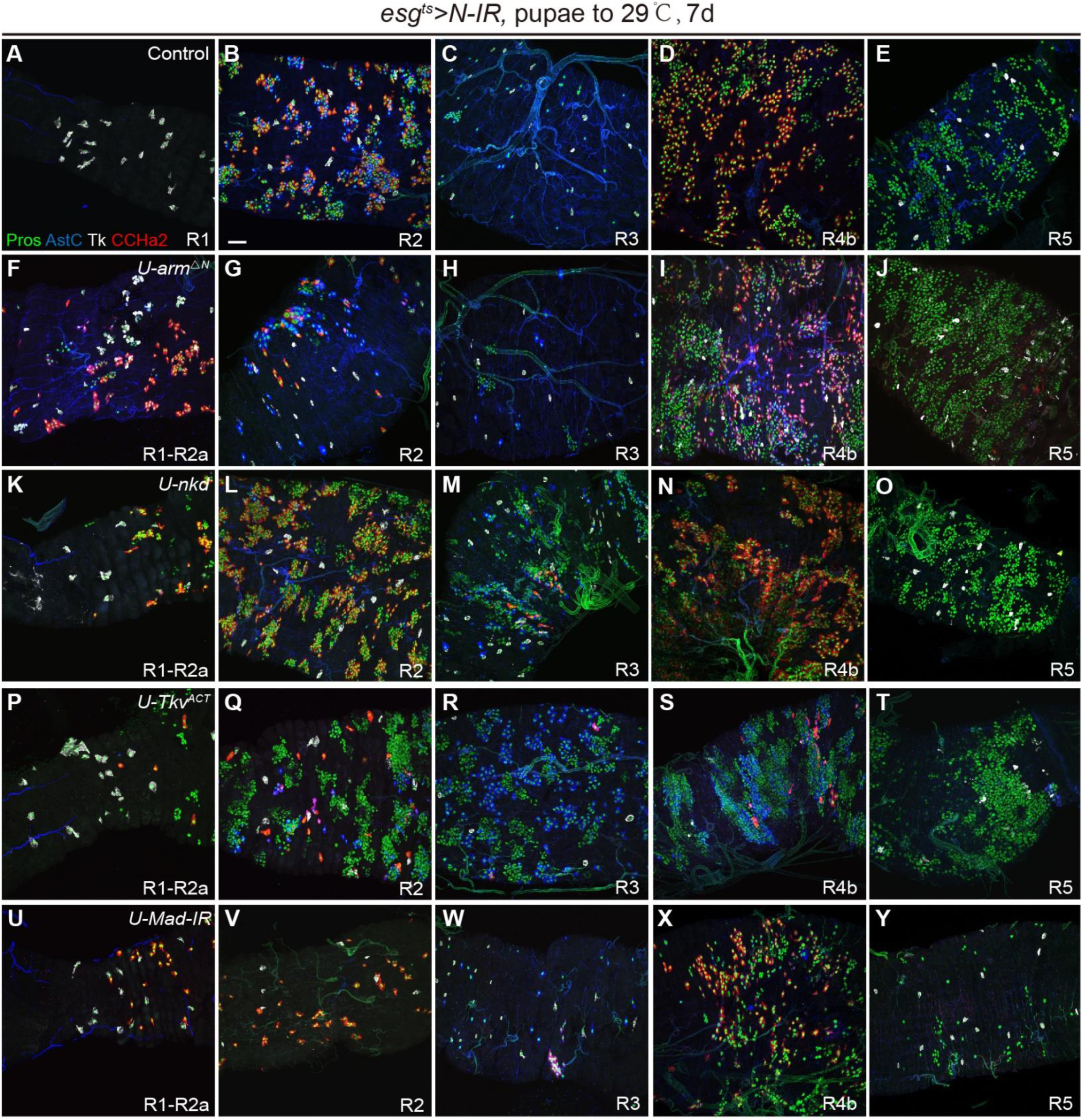
Manipulating WNT and BMP signaling with the loss of Notch in adult ISCs. (A-Y) Midguts stained with anti-Pros (green), anti-AstC (blue), anti-Tk (white) and anti-CCHa2 (red) in the following genotypes of flies driven by *esg-Gal4^ts^*with *UAS-N-IR* for 7 days since 48h APF: Control (A-E), *U-arm^S^*^10^ (F-J), *U-nkd* (K-O), *UAS-Tkv^ACT^* (P-T) and *UAS-Mad-IR* (U-Y). Scale bar represents 25 μm.

## References

1. Li, H., and Jasper, H. (2016). Gastrointestinal stem cells in health and disease: from flies to humans. Disease Models & Mechanisms 9, 487–499. 10.1242/dmm.024232.

2. Zwick, R.K., Kasparek, P., Palikuqi, B., Viragova, S., Weichselbaum, L., McGinnis, C.S., McKinley, K.L., Rathnayake, A., Vaka, D., Nguyen, V., et al. (2024). Epithelial zonation along the mouse and human small intestine defines five discrete metabolic domains. Nature Cell Biology 26, 250–262. 10.1038/s41556-023-01337-z.

3. Mattila, J., Viitanen, A., Fabris, G., Strutynska, T., Korzelius, J., and Hietakangas, V. (2024). Stem cell mTOR signaling directs region-specific cell fate decisions during intestinal nutrient adaptation. Science Advances 10. 10.1126/sciadv.adi2671.

4. Thompson, C.A., DeLaForest, A., and Battle, M.A. (2018). Patterning the gastrointestinal epithelium to confer regional-specific functions. Developmental Biology 435, 97–108. 10.1016/j.ydbio.2018.01.006.

5. Miguel-Aliaga, I., Jasper, H., and Lemaitre, B. (2018). Anatomy and Physiology of the Digestive Tract of Drosophila melanogaster. Genetics 210, 357–396. 10.1534/genetics.118.300224.

6. Guo, X., Lv, J., and Xi, R. (2021). The specification and function of enteroendocrine cells in Drosophila and mammals: a comparative review. The FEBS Journal 289, 4773–4796. 10.1111/febs.16067.

7. Gribble, F.M., and Reimann, F. (2019). Function and mechanisms of enteroendocrine cells and gut hormones in metabolism. Nat Rev Endocrinol 15, 226–237. 10.1038/s41574-019-0168-8.

8. Haber, A.L., Biton, M., Rogel, N., Herbst, R.H., Shekhar, K., Smillie, C., Burgin, G., Delorey, T.M., Howitt, M.R., Katz, Y., et al. (2017). A single-cell survey of the small intestinal epithelium. Nature 551, 333–339. 10.1038/nature24489.

9. Beumer, J., Puschhof, J., Bauzá-Martinez, J., Martínez-Silgado, A., Elmentaite, R., James, K.R., Ross, A., Hendriks, D., Artegiani, B., Busslinger, G.A., et al. (2020). High-Resolution mRNA and Secretome Atlas of Human Enteroendocrine Cells. Cell 181, 1291–1306.e1219. 10.1016/j.cell.2020.04.036.

10. Gehart, H., van Es, J.H., Hamer, K., Beumer, J., Kretzschmar, K., Dekkers, J.F., Rios, A., and Clevers, H. (2019). Identification of Enteroendocrine Regulators by Real-Time Single-Cell Differentiation Mapping. Cell 176, 1158–1173 e1116. 10.1016/j.cell.2018.12.029.

11. Grün, D., Lyubimova, A., Kester, L., Wiebrands, K., Basak, O., Sasaki, N., Clevers, H., and van Oudenaarden, A. (2015). Single-cell messenger RNA sequencing reveals rare intestinal cell types. Nature 525, 251–255. 10.1038/nature14966.

12. Guo, X., Yin, C., Yang, F., Zhang, Y., Huang, H., Wang, J., Deng, B., Cai, T., Rao, Y., and Xi, R. (2019). The Cellular Diversity and Transcription Factor Code of Drosophila Enteroendocrine Cells. Cell Rep 29, 4172–4185 e4175. 10.1016/j.celrep.2019.11.048.

13. Martin, A.M., Sun, E.W., and Keating, D.J. (2019). Mechanisms controlling hormone secretion in human gut and its relevance to metabolism. J Endocrinol 244, R1–R15. 10.1530/JOE-19-0399.

14. Yu, C.D., Xu, Q.J., and Chang, R.B. (2020). Vagal sensory neurons and gut-brain signaling. Curr Opin Neurobiol 62, 133–140. 10.1016/j.conb.2020.03.006.

15. Drucker, D.J. (2018). Mechanisms of Action and Therapeutic Application of Glucagon-like Peptide-1. Cell Metab 27, 740–756. 10.1016/j.cmet.2018.03.001.

16. Knudsen, L.B., and Lau, J. (2019). The Discovery and Development of Liraglutide and Semaglutide. Front Endocrinol (Lausanne) 10, 155. 10.3389/fendo.2019.00155.

17. Zwick, R.K., Ohlstein, B., and Klein, O.D. (2019). Intestinal renewal across the animal kingdom: comparing stem cell activity in mouse and Drosophila. American Journal of Physiology-Gastrointestinal and Liver Physiology 316, G313–G322. 10.1152/ajpgi.00353.2018.

18. Zhou, X., Ding, G., Li, J., Xiang, X., Rushworth, E., and Song, W. (2020). Physiological and Pathological Regulation of Peripheral Metabolism by Gut-Peptide Hormones in Drosophila. Front Physiol 11, 577717. 10.3389/fphys.2020.577717.

19. Buchon, N., Osman, D., David, F.P., Fang, H.Y., Boquete, J.P., Deplancke, B., and Lemaitre, B. (2013). Morphological and molecular characterization of adult midgut compartmentalization in Drosophila. Cell Rep 3, 1725–1738. 10.1016/j.celrep.2013.04.001.

20. Marianes, A., and Spradling, A.C. (2013). Physiological and stem cell compartmentalization within the Drosophila midgut. eLife 2. 10.7554/eLife.00886.

21. Ohlstein, B., and Spradling, A. (2006). The adult Drosophila posterior midgut is maintained by pluripotent stem cells. Nature 439, 470–474. 10.1038/nature04333.

22. Micchelli, C.A., and Perrimon, N. (2006). Evidence that stem cells reside in the adult Drosophila midgut epithelium. Nature 439, 475–479. 10.1038/nature04371.

23. Chen, J., Xu, N., Wang, C., Huang, P., Huang, H., Jin, Z., Yu, Z., Cai, T., Jiao, R., and Xi, R. (2018). Transient Scute activation via a self-stimulatory loop directs enteroendocrine cell pair specification from self-renewing intestinal stem cells. Nat Cell Biol 20, 152–161. 10.1038/s41556-017-0020-0.

24. Wu, S., Yang, Y., Tang, R., Zhang, S., Qin, P., Lin, R., Rafel, N., Lucchetta, E.M., Ohlstein, B., and Guo, Z. (2023). Apical-basal polarity precisely determines intestinal stem cell number by regulating Prospero threshold. Cell Reports 42. 10.1016/j.celrep.2023.112093.

25. Hung, R.J., Hu, Y., Kirchner, R., Liu, Y., Xu, C., Comjean, A., Tattikota, S.G., Li, F., Song, W., Ho Sui, S., and Perrimon, N. (2020). A cell atlas of the adult Drosophila midgut. Proc Natl Acad Sci U S A 117, 1514–1523. 10.1073/pnas.1916820117.

26. Chen, J., Kim, S.M., and Kwon, J.Y. (2016). A Systematic Analysis of Drosophila Regulatory Peptide Expression in Enteroendocrine Cells. Mol Cells 39, 358–366. 10.14348/molcells.2016.0014.

27. Jang, S., Chen, J., Choi, J., Lim, S.Y., Song, H., Choi, H., Kwon, H.W., Choi, M.S., and Kwon, J.Y. (2021). Spatiotemporal organization of enteroendocrine peptide expression in Drosophila. J Neurogenet 35, 387–398. 10.1080/01677063.2021.1989425.

28. Okamoto, N., and Watanabe, A. (2022). Interorgan communication through peripherally derived peptide hormones in Drosophila. Fly (Austin) 16, 152–176. 10.1080/19336934.2022.2061834.

29. Veenstra, J.A., Agricola, H.J., and Sellami, A. (2008). Regulatory peptides in fruit fly midgut. Cell Tissue Res 334, 499–516. 10.1007/s00441-008-0708-3.

30. Veenstra, J.A., and Ida, T. (2014). More Drosophila enteroendocrine peptides: Orcokinin B and the CCHamides 1 and 2. Cell Tissue Res 357, 607–621. 10.1007/s00441-014-1880-2.

31. Yoshinari, Y., Kosakamoto, H., Kamiyama, T., Hoshino, R., Matsuoka, R., Kondo, S., Tanimoto, H., Nakamura, A., Obata, F., and Niwa, R. (2021). The sugar-responsive enteroendocrine neuropeptide F regulates lipid metabolism through glucagon-like and insulin-like hormones in Drosophila melanogaster. Nat Commun 12, 4818. 10.1038/s41467-021-25146-w.

32. Kubrak, O., Koyama, T., Ahrentlov, N., Jensen, L., Malita, A., Naseem, M.T., Lassen, M., Nagy, S., Texada, M.J., Halberg, K.V., and Rewitz, K. (2022). The gut hormone Allatostatin C/Somatostatin regulates food intake and metabolic homeostasis under nutrient stress. Nat Commun 13, 692. 10.1038/s41467-022-28268-x.

33. Li, H., Qi, Y., and Jasper, H. (2013). Dpp signaling determines regional stem cell identity in the regenerating adult Drosophila gastrointestinal tract. Cell Rep 4, 10–18. 10.1016/j.celrep.2013.05.040.

34. Driver, I., and Ohlstein, B. (2014). Specification of regional intestinal stem cell identity during Drosophila metamorphosis. Development 141, 1848–1856. 10.1242/dev.104018.

35. Beehler-Evans, R., and Micchelli, C.A. (2015). Generation of enteroendocrine cell diversity in midgut stem cell lineages. Development 142, 654–664. 10.1242/dev.114959.

36. Takashima, S., Adams, K.L., Ortiz, P.A., Ying, C.T., Moridzadeh, R., Younossi-Hartenstein, A., and Hartenstein, V. (2011). Development of the Drosophila entero-endocrine lineage and its specification by the Notch signaling pathway. Dev Biol 353, 161–172. 10.1016/j.ydbio.2011.01.039.

37. Micchelli, C.A., Sudmeier, L., Perrimon, N., Tang, S., and Beehler-Evans, R. (2011). Identification of adult midgut precursors in Drosophila. Gene Expr Patterns 11, 12–21. 10.1016/j.gep.2010.08.005.

38. Guo, Z., and Ohlstein, B. (2015). Bidirectional Notch signaling regulates Drosophila intestinal stem cell multipotency. Science 350. 10.1126/science.aab0988.

39. Amcheslavsky, A., Jiang, J., and Ip, Y.T. (2009). Tissue damage-induced intestinal stem cell division in Drosophila. Cell Stem Cell 4, 49–61. 10.1016/j.stem.2008.10.016.

40. Yin, C., and Xi, R. (2018). A Phyllopod-Mediated Feedback Loop Promotes Intestinal Stem Cell Enteroendocrine Commitment in Drosophila. Stem Cell Reports 10, 43–57. 10.1016/j.stemcr.2017.11.014.

41. Guo, X., Zhang, Y., Huang, H., and Xi, R. (2022). A hierarchical transcription factor cascade regulates enteroendocrine cell diversity and plasticity in Drosophila. Nat Commun 13, 6525. 10.1038/s41467-022-34270-0.

42. Lee, T., and Luo, L. (1999). Mosaic Analysis with a Repressible Cell Marker for Studies of Gene Function in Neuronal Morphogenesis. Neuron 22, 451–461. 10.1016/s0896-6273(00)80701-1.

43. Tian, A., Benchabane, H., Wang, Z., and Ahmed, Y. (2016). Regulation of Stem Cell Proliferation and Cell Fate Specification by Wingless/Wnt Signaling Gradients Enriched at Adult Intestinal Compartment Boundaries. PLoS Genet 12, e1005822. 10.1371/journal.pgen.1005822.

44. Tian, A., Duwadi, D., Benchabane, H., and Ahmed, Y. (2019). Essential long-range action of Wingless/Wnt in adult intestinal compartmentalization. PLoS Genet 15, e1008111. 10.1371/journal.pgen.1008111.

45. Lin, G., Xu, N., and Xi, R. (2008). Paracrine Wingless signalling controls self-renewal of Drosophila intestinal stem cells. Nature 455, 1119–1123. 10.1038/nature07329.

46. Wang, C., Zhao, R., Huang, P., Yang, F., Quan, Z., Xu, N., and Xi, R. (2013). APC loss-induced intestinal tumorigenesis in Drosophila: Roles of Ras in Wnt signaling activation and tumor progression. Dev Biol 378, 122–140. 10.1016/j.ydbio.2013.03.020.

47. Guo, Z., Driver, I., and Ohlstein, B. (2013). Injury-induced BMP signaling negatively regulates Drosophila midgut homeostasis. J Cell Biol 201, 945–961. 10.1083/jcb.201302049.

48. Mehrotra, S., Bansal, P., Oli, N., Pillai, S.J., and Galande, S. (2020). Defective Proventriculus Regulates Cell Specification in the Gastric Region of Drosophila Intestine. Frontiers in Physiology 11. 10.3389/fphys.2020.00711.

49. Guo, X., Wang, C., Zhang, Y., Wei, R., and Xi, R. (2024). Cell-fate conversion of intestinal cells in adult Drosophila midgut by depleting a single transcription factor. Nat Commun 15, 2656. 10.1038/s41467-024-46956-8.

50. Dutta, D., Dobson, Adam J., Houtz, Philip L., Gläßer, C., Revah, J., Korzelius, J., Patel, P.H., Edgar, Bruce A., and Buchon, N. (2015). Regional Cell-Specific Transcriptome Mapping Reveals Regulatory Complexity in the Adult Drosophila Midgut. Cell Reports 12, 346–358. 10.1016/j.celrep.2015.06.009.

51. Zhou, J., Florescu, S., Boettcher, A.-L., Luo, L., Dutta, D., Kerr, G., Cai, Y., Edgar, B.A., and Boutros, M. (2015). Dpp/Gbb signaling is required for normal intestinal regeneration during infection. Developmental Biology 399, 189–203. 10.1016/j.ydbio.2014.12.017.

52. Beumer, J., Artegiani, B., Post, Y., Reimann, F., Gribble, F., Nguyen, T.N., Zeng, H., Van den Born, M., Van Es, J.H., and Clevers, H. (2018). Enteroendocrine cells switch hormone expression along the crypt-to-villus BMP signalling gradient. Nat Cell Biol 20, 909–916. 10.1038/s41556-018-0143-y.

53. Fujii, M., Matano, M., Toshimitsu, K., Takano, A., Mikami, Y., Nishikori, S., Sugimoto, S., and Sato, T. (2018). Human Intestinal Organoids Maintain Self-Renewal Capacity and Cellular Diversity in Niche-Inspired Culture Condition. Cell Stem Cell 23, 787–793 e786. 10.1016/j.stem.2018.11.016.

54. Middendorp, S., Schneeberger, K., Wiegerinck, C.L., Mokry, M., Akkerman, R.D., van Wijngaarden, S., Clevers, H., and Nieuwenhuis, E.E. (2014). Adult stem cells in the small intestine are intrinsically programmed with their location-specific function. Stem Cells 32, 1083–1091. 10.1002/stem.1655.

55. Fukuda, M., Mizutani, T., Mochizuki, W., Matsumoto, T., Nozaki, K., Sakamaki, Y., Ichinose, S., Okada, Y., Tanaka, T., Watanabe, M., and Nakamura, T. (2014). Small intestinal stem cell identity is maintained with functional Paneth cells in heterotopically grafted epithelium onto the colon. Genes Dev 28, 1752–1757. 10.1101/gad.245233.114.

56. Brand, A.H., and Perrimon, N. (1993). Targeted gene expression as a means of altering cell fates and generating dominant phenotypes. Development 118, 401–415.

57. McGuire, S.E., Mao, Z., and Davis, R.L. (2004). Spatiotemporal gene expression targeting with the TARGET and gene-switch systems in Drosophila. Sci STKE 2004, pl6.

58. Lai, S.L., and Lee, T. (2006). Genetic mosaic with dual binary transcriptional systems in Drosophila. Nat Neurosci 9, 703–709. 10.1038/nn1681.

59. Veenstra, J.A., and Ida, T. (2014). More Drosophila enteroendocrine peptides: Orcokinin B and the CCHamides 1 and 2. Cell and Tissue Research 357, 607–621. 10.1007/s00441-014-1880-2.

60. Kunst, M., Hughes, M.E., Raccuglia, D., Felix, M., Li, M., Barnett, G., Duah, J., and Nitabach, M.N. (2014). Calcitonin gene-related peptide neurons mediate sleep-specific circadian output in Drosophila. Curr Biol 24, 2652–2664. 10.1016/j.cub.2014.09.077.

61. Reichwald, K., Unnithan, G.C., Davis, N.T., Agricola, H., and Feyereisen, R. (1994). Expression of the allatostatin gene in endocrine cells of the cockroach midgut. Proc Natl Acad Sci U S A 91, 11894–11898. 10.1073/pnas.91.25.11894.

62. Lee, S.H., Cho, E., Yoon, S.E., Kim, Y., and Kim, E.Y. (2021). Metabolic control of daily locomotor activity mediated by tachykinin in Drosophila. Commun Biol 4, 693. 10.1038/s42003-021-02219-6.

